# Watch-breaker: establishment of a microwell array-based miniaturized thymic organoid model suitable for high throughput applications

**DOI:** 10.1101/2024.09.23.614441

**Authors:** Viktoria Major, Sam Palmer, Paul Rouse, Timothy Henderson, Tania Hübscher, Joanna Sweetman, Jan Morys, Andrea Bacon, An Chengrui, Qiu Guiyun, Yu Wang, Andrea Corsinotti, Justyna Cholewa-Waclaw, S. Jon Chapman, Matthias P. Lütolf, Graham Anderson, C. Clare Blackburn

**Affiliations:** Institute of Regeneration and Repair, Centre for Regenerative Medicine, University of Edinburgh, 5 Little France Drive, Edinburgh EH16 4UU, UK; Institute for Stem Cell Research, School of Biological Sciences, University of Edinburgh.; Mathematical Institute, University of Oxford, Woodstock Road, Oxford, OX2 6GG, UK; École Polytechnique Fédérale de Lausanne, EPFL SV IBI-SV UPLUT, AI 1208 (Bâtiment AI), Station 15, CH-1015 Lausanne.; Institute of Immunology and Immunotherapy, University of Birmingham, B15 2TT, UK

**Keywords:** Thymus, epithelial cells, organoid, high throughput, in vitro T cell development, microphysiological

## Abstract

T-cell development depends critically on the thymic stroma, in particular the diverse array of functionally distinct thymic epithelial cell (TEC) types. However, a robust *in vitro* thymus model mimicking the native thymus and compatible with medium/high-throughput analyses is currently lacking. Here, we demonstrate a novel high-density microwell array-based miniaturized thymus organoid (mTO) model, that supports T-cell commitment and development, possesses key organizational characteristics of the native thymus and is compatible with live-imaging and medium/high-throughput applications. We establish the minimum cellular input required for functional mTO and show that mTO TEC phenotype and complexity closely mirrors the native thymus. Finally, we use mTO to probe the role of fetal thymic mesenchyme, revealing a requirement beyond maintenance of *Foxn1* in differentiation/maintenance of mature TEC subpopulations. Collectively, mTO present a new *in vitro* model of the native thymus adaptable to medium/high-throughput applications and validated for exploration of thymus- and thymus organoid-biology.

## Introduction

The thymus is required for T-cell development. Obligate dynamic interactions with thymic stromal cells regulate haematopoietic progenitor cell (HPC) colonisation, commitment of HPCs to the T-cell lineage, and subsequent progression through the T-cell differentiation and selection pathways^1-4^. The thymic stroma comprises epithelial, mesenchymal, vascular and haematopoietic components^5-7^, among which thymic epithelial cells (TEC) are critical for the organ’s specialist functions^4^. The functionally distinct cortical (c) and medullary (m) TEC sub-lineages define the two main thymic compartments each contain several sub-types^8-13^. Broadly, cTEC regulate T-cell lineage commitment, differentiation and positive selection while mTEC regulate central tolerance induction^4,14-19^; these distinct functions depend on unique, sub-lineage specific gene expression programmes^4,8,20-23^. TEC also regulate the production of ψ8 T-cells and non-conventional innate subsets, including iNKTs.

The current strong interest in thymus biology and T-cell development is underpinned by three main factors: (i) The thymus undergoes a stereotypical age-related involution, resulting in age-related decline in new naïve T-cell output that contributes to the increased risk of cancer and infection with age, and adversely affecting recovery of immune function after cytoablative therapies and chronic infection^24-26^; (ii) Some immunodeficiencies are caused by failure of thymus function or by early life surgical thymectomy; and (iii) T-cells have recently been harnessed for successful cancer immunotherapies. However, thymus transplants currently rely on neonatal thymus tissue obtained as surgical discard, function only transiently and are frequently associated with autoimmunity^27^, and neither clinically useful thymus regeneration nor the generation of bespoke T-cell repertoires *in vitro* have yet been achieved^26^.

An *in vitro* experimental platform compatible with medium/high-throughput mechanistic interrogation of physiological thymus functions would enable progress in all the above areas. Existing models however each have substantive limitations. While fetal thymic organ culture (FTOC) and reaggregate fetal thymic organ culture (RFTOC)^28^ largely recapitulate physiological function, they depend on use of fetal thymic cells - precluding medium/high throughput approaches - and neither supports *in vitro* genetic manipulation of thymic stroma. Cell-line models based on the mouse bone marrow-derived cell lines OP9 or MS5 transduced with the Notch ligands Delta like 1 (DL1) or Delta like 4 (DLL4) partially support T-cell development^29-33^. However, these models lack cTEC- and mTEC-specific molecular machineries required for physiological positive and negative selection. Additionally, as OP9 and MS5 are not related to TEC they do not support studies investigating thymic stromal cell function. Recently, progress has been made in direct reprogramming of unrelated cell types into TEC^34^, directed differentiation of mouse and human pluripotent stem cells (PSC) into thymic epithelial progenitor cells (TEPC)^35-38^, and using cultured mouse and human TEC as the basis of transplanted or *in vitro* thymic organoids^39-43^. However, although promising for further development, these models largely remain poorly characterised relative to *ex vivo* TEC, RFTOC and FTOC. Thus, improved models are required.

Miniaturized organoids that mimic physiological function have recently been established for some organs and tissues. Thus, we set out to test whether this approach could be applied to the thymus. Here, we demonstrate that miniaturized thymic organoids (mTO) able to mediate thymopoiesis can be established from *ex vivo* thymus tissue. We define the minimum cellular inputs required to generate functional mTO and demonstrate the presence within these organoids of discrete cTEC and mTEC regions including AIRE^+^ mTEC. Via single cell transcriptomics we establish that most if not all normal intrathymic stromal cell-types are present within mTO and reveal likely pathways through which fetal thymic mesenchyme (FTM) impacts TEC development and function. This novel microphysiological thymic organoid system reduces variability and input cell numbers compared to other approaches, while retaining cTEC:mTEC compartmentalisation and cellular diversity. Importantly, it is suitable for live/timelapse imaging and medium/high throughput screening applications, representing a step-change for interrogation of thymic stromal biology.

## Results

The essential features of any functional thymic organoid are: (i) ability to support T-cell lineage commitment from HPC; (ii) ability to support subsequent thymocyte differentiation to generate a pool of CD4^+^ and CD8^+^ T-cells exhibiting T-cell receptor (TCR) diversity; (iii) regions of cTEC, that express DLL4 (HPC commitment to the T-cell lineage^44,45^); CXCL12 (β-selection of developing T-cells^46^); β5t (cTEC -specific proteasome subunit that generates the optimum peptide repertoire for positive selection of CD8^+^ T-cells^47^); Cathepsin L (cTEC-specific protease required for optimum positive selection of CD4^+^ T-cells^48^); MHC Class I and MHC Class II (positive selection of CD8^+^ and CD4^+^ T-cells respectively); and (iv) expression by the TEC of cytokines needed to drive proliferation and differentiation of the developing T-cells (e.g. Flt3L, CXCL12, IL-7). An organoid containing these features should generate a repertoire of CD4^+^ and CD8^+^ T-cells that have been positively-selected on the peptide repertoires used in physiological positive selection and so will be responsive to antigenic peptides in the correct affinity range. Thus, establishment of miniaturized thymic organoids (mTO) that met criteria (i-iv) above was the minimal goal of our study. The presence of medullary regions within thymic organoids, evidenced by the presence of TEC expressing cytokeratin 14 (K14), Autoimmune Regulator (AIRE), FEZF2 and a range of AIRE-dependent and AIRE-independent tissue restricted antigens (TRAs) would promote central tolerance induction in the positively-selected repertoire and would also be desirable for some applications.

### mTO form in microwells and mediate thymopoiesis

To test whether thymic reaggregates could be established in microwell plates we seeded defined numbers of dissociated fetal thymus cells from a range of developmental stages into single wells of Gri3D^®^ 96 well plates (SUN Biosciences) in which each well contains a U-bottomed microwell-printed PEG-based hydrogel insert (Figure 1A). Preliminary testing established that seeding cells from two thymic lobes (i.e. one embryo equivalent) per well, in which each well contained thirty-one 800µm diameter microwells, performed better than any of the other conditions tested (different input cell numbers and microwell diameters were tested; not shown).

**Figure 1:**
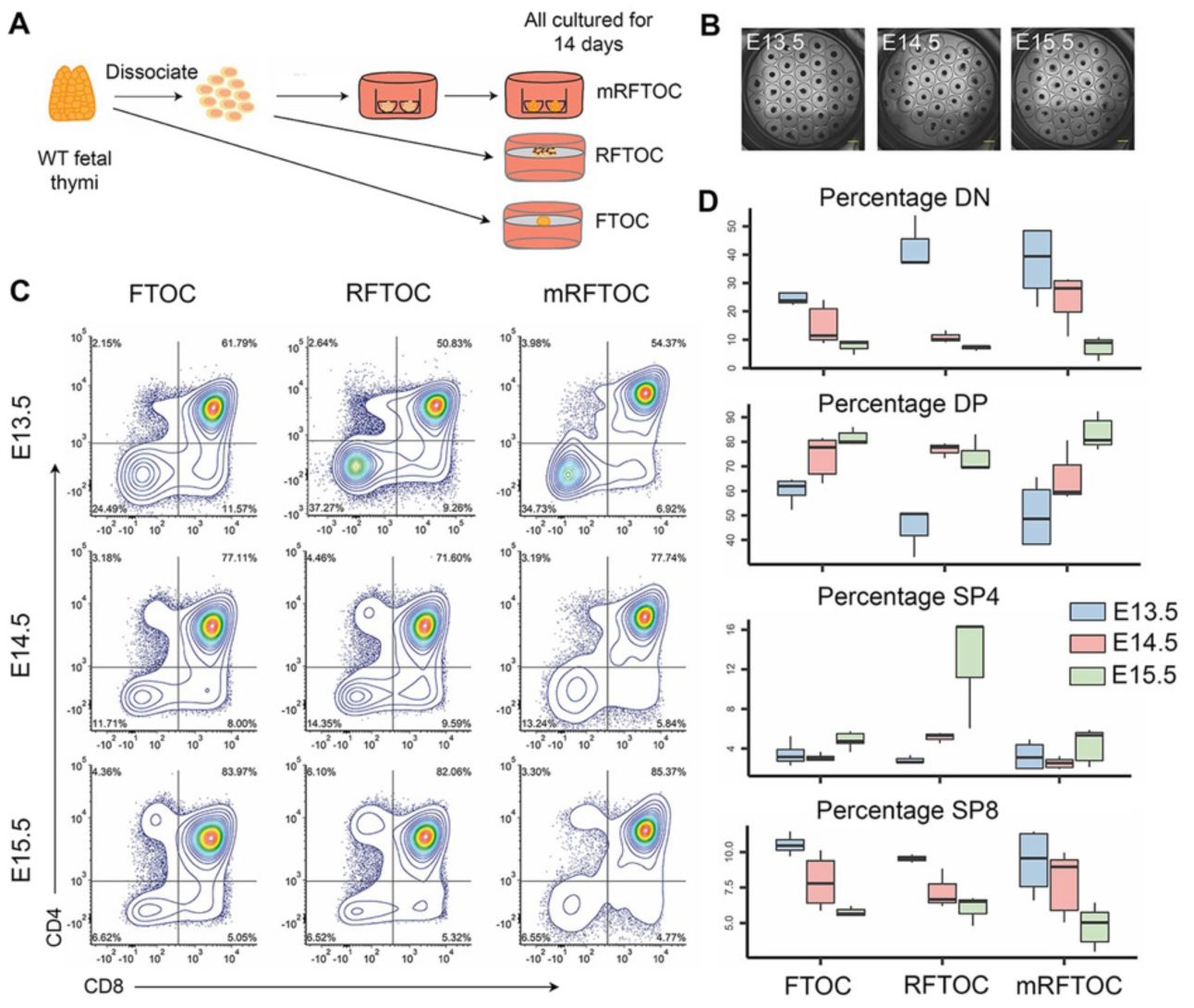
Thymopoiesis in unsorted mTO, FTOC and RFTOC. (A) Schematic depicting creation of FTOC, RFTOC and unsorted mTO. (B) Bright field image of E13.5 (top), E14.5 (middle) and E15.5 (bottom) unsorted mTO seven days after seeding, each micro-well contains a reaggregate. (C, D) Thymocyte subset analysis after seven days culture for the conditions and input tissue ages shown. (C) Graphs show percentage of DN, DP, SP4 and SP8 thymocyte populations. (D) Representative CD4 versus CD8 plots for FTOC (left), RFTOC (centre) and unsorted mTO (right) from the input tissue ages shown. (C,D) Data shown are for live lineage negative cells (lin=CD11b, CD11c, Gr-1, Nk1.1, B220, EpCAM and Ter119). N=3 independent biological replicates for each experiment. (B) Scale bars = 500μm.

We then analysed the effect of developmental stage on reaggregate formation, choosing embryonic day 13.5 (E13.5), E14.5 and E15.5 fetal thymus cells as our input populations (these stages exhibit progressively increasing levels of TEC differentiation and complexity^49-51^). Dissociated-but-unsorted fetal thymus cells from each of the stages formed cellular aggregates in most if not all microwells by 24 hours after seeding, and formed clear spheroid structures by 48 hours. The spheroids persisted for at least fourteen days in culture (the latest timepoint analyzed; Figure 1B), at which timepoint double negative (DN; CD8^-^CD4^-^), double positive (DP; CD8^+^CD4^+^), CD4^+^ single positive (SP4) and CD8^+^ single positive (SP8) cells were present in all conditions. Thymocyte subset profiles were comparable to FTOC and standard RFTOC controls (Figure 1C, D; note that each standard RFTOC was formed from six thymic lobes; at E14.5 each lobe contains approximately 1x10^5^ total cells). E14.5 mTO performed very similarly to FTOC and RFTOC in terms of proportions of DP, SP4 and SP8 present, and E14.5 thymi were thus used for all further analyses.

### Defining the minimal cellular inputs for functional mTO formation

We next determined the minimal cellular inputs for establishment of functional mTO that met criteria (i-iv) above, asking whether and in what proportion each of the major thymic stromal populations was required (Figure 2A). We called this experiment ‘Watch-breaker’. E14.5 thymi were microdissected, dissociated to single cell suspensions and separated into TEC, FTM, haematopoietic (CD45^+^; at E14.5 primarily DN1-3, see Figure S1) and endothelial cell (EC) populations by flow cytometric sorting. At E14.5, of live cells, sixty percent were thymocytes, thirty percent TEC, eleven percent FTM and less than one percent ECs (Figure S1), with around four percent of TEC being UEA1^+^ mTEC progenitors^50^.

**Figure 2:**
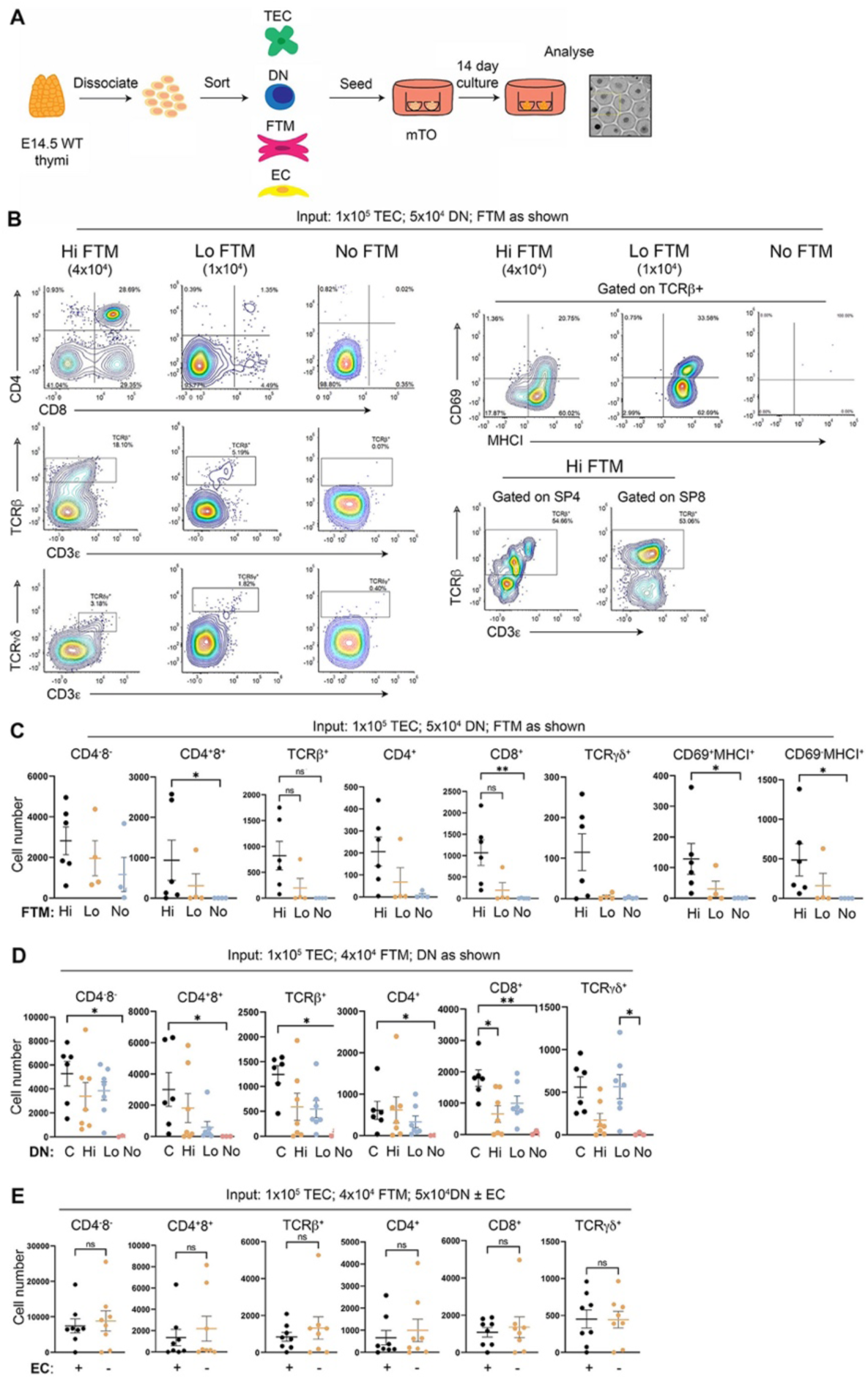
Establishment of the minimum cellular inputs for functional mTO. (A) Schematic of experimental design. (B, C) mTO were generated with a constant number of TEC and DN and varying numbers of FTM cells: “Hi FTM” – 1x10^5^ TEC, 5x10^4^ DN, 4x10^4^ FTM per well (physiological FTM:TEC ratio); “Lo FTM” - 1x10^5^ TEC, 5x10^4^ DN, 1.5x10^4^ FTM; and “No-FTM” - 1x10^5^ TEC, 5x10^4^ DN, 0 FTM. EC were not included. (B) Representative plots showing flow cytometric analysis of thymocyte subset distribution after 14 days’ culture, for the markers and conditions shown. (C) Absolute numbers of recovered thymocytes from (B). (D, E) mTO were generated with a constant number of TEC and FTM and varying numbers of DN cells (D), or with and without thymic EC (E): (D) All conditions – 1x10^5^ TEC, 4x10^4^ FTM; Control – 5x10^4^ DN; “Hi DN” - 2x10^5^ DN; “Lo DN” 1x10^4^ DN. (E) No EC control – 1x10^5^ TEC, 5x10^4^ DN, 4x10^4^ FTM; EC, as control with 1-2x10^4^ EC. Plots show absolute numbers of recovered thymocytes for each subset. (B, C) n=5, (D) n=8, (F) n=8, where n is an independent biological replicate performed on a separate day. Cell numbers shown are per well of a single 96 well. C, control. (B-E) Data shown are for live CD45^+^ lineage negative cells (lin=CD11b, CD11c, Gr-1, Nk1.1, B220, EpCAM and Ter119); CD69^+^MHCI^+^ and CD69^-^MHCI^+^ data shown are after subsequent gating on TCRβ^+^ cells. See also Figures S1-4.

We first tested the requirement for FTM and of altering the FTM:TEC ratio (Figure 2A-B), using three conditions: “Hi FTM” – 1x10^5^ TEC, 5x10^4^ DN, 4x10^4^ FTM per well (the physiological FTM:TEC ratio); “Lo FTM” - 1x10^5^ TEC, 5x10^4^ DN, 1.5x10^4^ FTM; and “No-FTM” - 1x10^5^ TEC, 5x10^4^ DN, 0 FTM. EC were not included. The ‘Hi’ and ‘Lo’ FTM conditions both sustained thymopoiesis, evidenced by the presence of DP, SP4 and SP8 populations after fourteen days. TCRβ^hi^ ‘SP4 and SP8 and TCRψ8^+^ thymocytes were present in both ‘Hi’ and ‘Lo’ FTM conditions; the SP cells had undergone positive selection and subsequent maturation, evidenced by the presence of CD69^+^MHCI^+^ (M1) and CD69^-^MHCI^+^ (M2) populations (Figure 2B,C, Figure S2). In the absence of FTM, thymocyte development did not proceed beyond the DN stage (Figure 2B,C, Figure S2). The ‘Hi FTM’ condition resulted in greatly increased output cell numbers compared to ‘Lo FTM’ (see Figure 2C) and was thus adopted for all further experiments.

We then optimized the input thymocyte numbers, by generating mTO in which TEC and FTM numbers were as for the ‘Hi FTM’ condition (1x10^5^ TEC, 4x10^4^ FTM) while DN thymocyte number was varied (“Hi DN” - 2x10^5^ DN; “Control” 5x10^4^ DN (i.e. as in “Hi FTM”); “Lo DN” 1x10^4^ DN; “No DN”, 0 DN cells). As expected, no thymocytes were recovered from the “No DN” condition. The “Hi DN”, “Lo DN” and “Control” mTO all sustained thymopoiesis and produced SP4 and SP8 αβ and ψ8 T-cells (Figure 2D). The “Control” (i.e. “Hi FTM”) condition produced the highest number of all thymocyte subsets (Figure 2D) with better maturation through the CD69^+^MHCI^+^ and CD69^-^ MHCI^+^ stages (not shown), and also demonstrated less variability across the individual replicates than the other conditions. We therefore continued to use this condition for all further analysis. Notably, although the “Hi DN” condition represented the physiological E14.5 TEC:DN:FTM ratio it performed worse than the “Control” (i.e. “Hi FTM”) condition in which DN were proportionally lower, in terms of output cell numbers.

We also tested whether the addition of thymic EC to mTO would affect functionality. No significant differences were found between the ‘With EC’ and ‘Without EC’ conditions tested (1x10^5^ TEC, 5x10^4^ DN, 4x10^4^ FTM, ±1-2x10^4^ EC)(Figure 2E).

Finally, we tested whether increasing or decreasing the seeding cell number per well would affect mTO function and whether ‘pre-mixing’ of the different cell inputs affected mTO function compared to the sequential seeding approach used above. For this, we established mTO using the optimized “Hi FTM” ratios but with the total cell numbers set at 1.5x or 0.5x the 1x “Hi FTM” control. We also established mTO in which the seeding cells were either pre-mixed or seeded sequentially. The 1x Hi FTM and 0.5x Hi FTM conditions produced the highest number of cells per input CD45^+^ cell for each thymocyte subset tested, with the Hi FTM condition performing better in terms of numbers of SP4 and SP8 produced (s). Therefore, this was retained as the optimized condition for further work. No significant differences were found for sequential seeding versus pre-mixing, and therefore the pre-mixing protocol was adopted for subsequent experiments (Figure S4).

### mTO support thymopoiesis from uncommitted haematopoietic progenitors

The above data used differentiation of DN thymocytes to read out mTO function. Since some of the E14.5 DN input cells were at or beyond the CD44^+^CD25^+^ DN2 stage (i.e. had commenced T-lineage differentiation), we also tested whether mTO could mediate T-cell commitment. For this, we isolated E14.5 fetal liver lymphoid-primed multipotent progenitors (LMPP) and substituted these for the DN cells such that the input cell numbers were 1x10^5^ TEC, 3.1x10^3^ LMPP, 4x10^4^ FTM, with the control being 1x10^5^ TEC, 5x10^4^ DN, 4x10^4^ FTM (i.e. Hi FTM). mTO supported robust thymopoiesis from LMPP, albeit with delayed differentiation kinetics compared to DN cells as evidenced by the low proportion of TCRβ^hi^ cells in the LMPP condition at the timepoint assayed (Figure 3A, B). Indeed, the number of DP, SP4, SP8 and ψ8 thymocytes generated per input cell was higher for LMPP than for DN (Figure 3B; input numbers of LMPP were 16-fold lower than DN).

**Figure 3:**
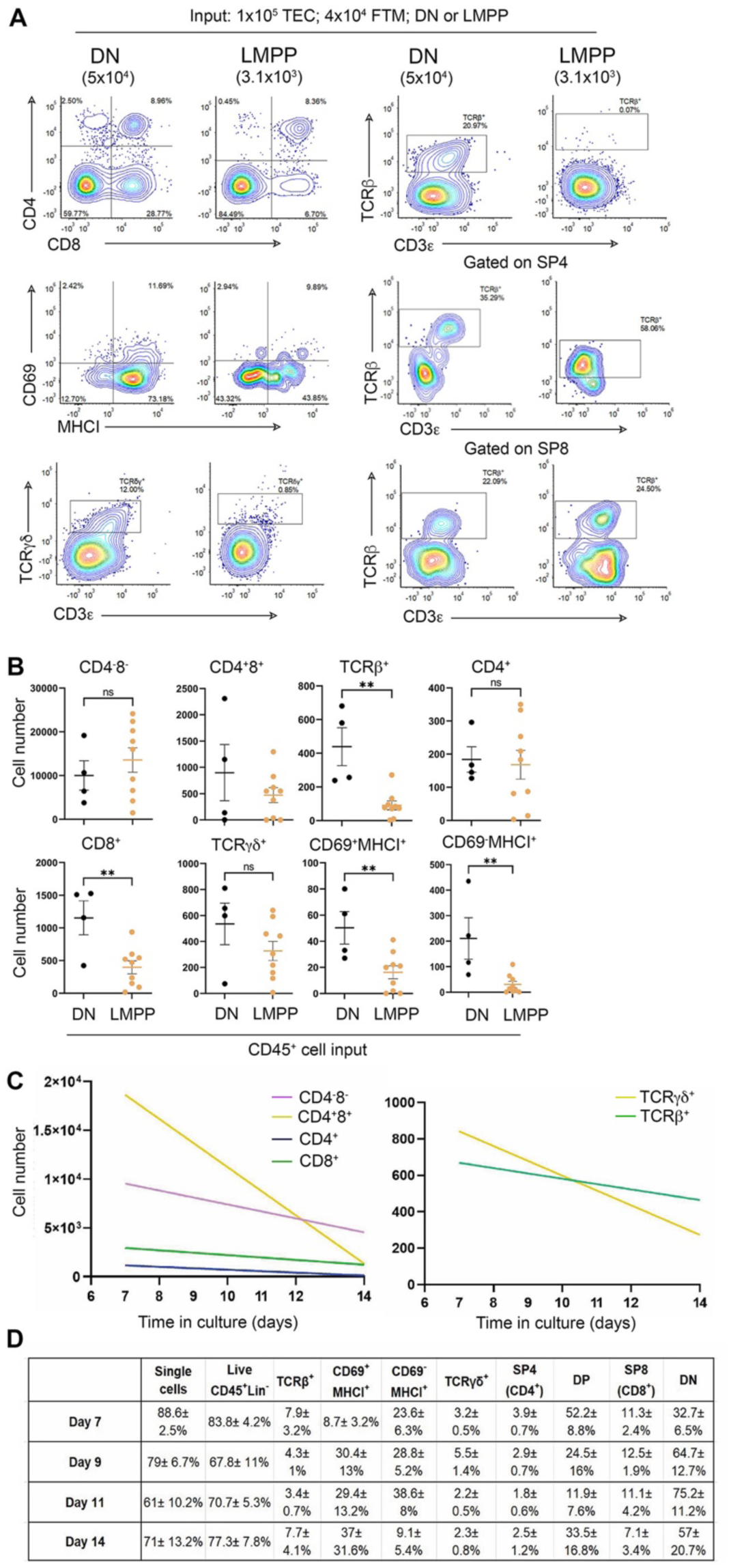
Thymopoiesis from uncommitted haematopoietic progenitors. (A, B) mTO were established from E14.5 thymic cells under control conditions (1x10^5^ TEC, 5x10^4^ DN, 4x10^4^ FTM) or with 3.1x10^3^ E14.5 fetal liver LMPP instead of DN input cells. (A) Plots show representative flow cytometric analysis for the markers shown, after 14 days’ culture for the conditions shown. (B) Absolute numbers of recovered thymocytes for each subset. (C,D) mTO established under control conditions (1x10^5^ TEC, 5x10^4^ DN, 4x10^4^ FTM) were analysed for thymocyte subset distribution at the timepoints shown. (C) Graphs show absolute cell numbers for the populations and timepoints shown. (D) Table shows percentages for each population among all live lin^-^ cells. (A-D) Each data point represents the cells harvested from one microwell. Graphs and table show mean± SEM. n=4 for each experiment, where n is an independent biological replicate performed on a separate day, with a total of 9 technical replicates across n=4. Statistical analysis was by unpaired t-test or Mann-Whitney rank test based on Shapiro-Wilk normality test. (A-D) Data shown are for live CD45^+^ lineage negative cells (lin=CD11b, CD11c, Gr-1, Nk1.1, B220, EpCAM and Ter119); CD69^+^MHCI^+^ and CD69^-^MHCI^+^ data shown are after subsequent gating on TCRβ^+^ cells. See also Figure S5.

### Kinetics of T-cell development within mTO

To analyse the kinetics of T-cell development within mTO, we also analysed “Hi FTM” mTO for thymocyte developmental progression and evidence of positive selection and SP maturation at range of timepoints: day 7, 9, 11 and 14 of culture. A striking downward trend in absolute DP numbers was evident as time in culture increased, from almost 2x10^4^ (day 7) to less than 5x10^3^ (day 14) (Figure 3C,D, Figure S5). The absolute number of recovered SP4 and SP8 thymocytes also decreased, though less dramatically than for DP (Figure 3C,D, Figure S5) and a decrease γδ T-cell numbers over time was also observed (Figure 3C,D, Figure S5). Representative overview images showed more consistent mTO morphology at Days 7, 9 and 11 compared to Day 14 (not shown). Overall, the 7-day culture period yielded peak numbers of developing T-cells and showed the lowest experiment-to-experiment variation for all populations tested, so was adopted for further analyses (Figure 3D, Figure S5; live CD45^+^ cells recovered: 7 day endpoint, 5.41x10^4^±2.38x10^4^ versus 14 day endpoint, 9.2x10^3^±1.58x10^3^).

Collectively, mTO established from input E14.5 thymus cells can support thymopoiesis from both DN cells and LMPP, with CD69 expression in DP evidencing positive selection; formation of optimally functional mTO requires FTM, but T-cell development within mTO does not require thymic endothelial cells; and a 7 day culture period results in maximal cell numbers while 12-14 days culture yields more TCRβ^hi^ cells.

### Segregation of cortex-like and medulla-like regions in mTO

We next assessed the organisation and identity of stromal cells within mTO (Figure 4). To analyse the distribution of TEC we established mTO from two reporter mouse strains, *Foxn1^GFP^* ^52^ and *Rank-Venus;Cxcl12dDsRed*^53-55^. FOXN1 is a master regulator of TEC differentiation required for development of all mature TEC subpopulations^49^ while RANK and CXCL12 mark mTEC and cTEC, respectively.

**Figure 4:**
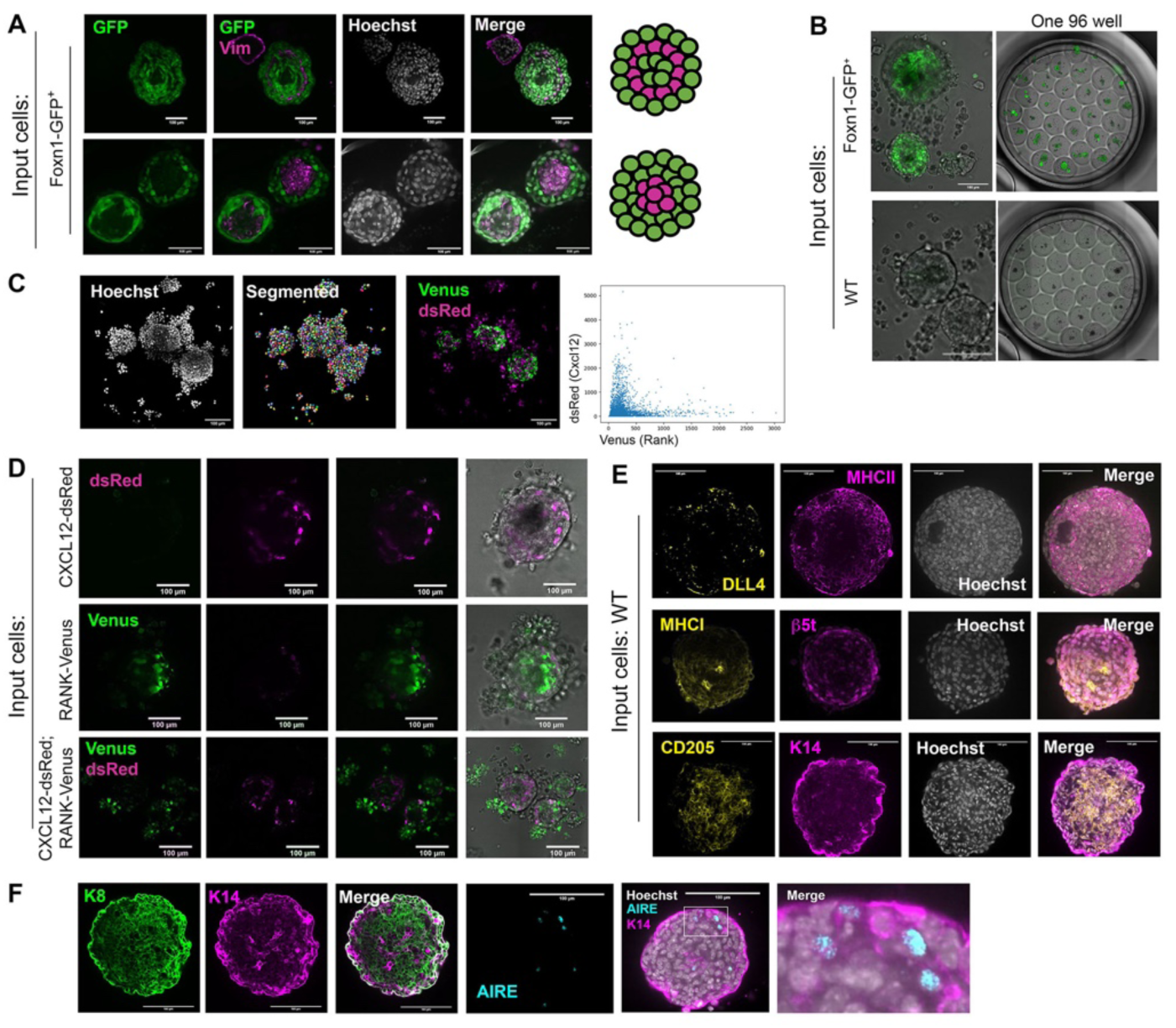
cTEC and mTEC regions are segregated in mTO. Representative images of individual mTOs established from E14.5 WT, Foxn1^GFP^, Cxcl12-dsRED, Rank-VENUS or Cxcl12-DsRed;Rank-Venus thymi and cultured for 7 days. All images shown are single optical sections taken from the centre of the mTOs, after live imaging at 40x magnification using an Opera Phenix. Scale bars,100µm. (A) Images are representative of Foxn1^GFP^ mTOs, showing direct GFP fluorescence (green) and staining for vimentin (magenta) and nuclei (Hoecsht, white). Right hand panels show schematic representation of different staining patterns observed in (A). (B-F) Images are representative of (B) Foxn1^GFP^ showing direct GFO fluorescence (green); (C, D) Cxcl12-dsRED, Rank-VENUS or Cxcl12-DsRed;Rank-Venus mTOs showing direct Venus (green) and dsRed (magenta) fluorescence, and merged channels with brightfield. Right hand panel in C shows dsRed and Venus intensity in individual cells as identified by segmentation in adjacent image; (E) WT mTOs showing β5t, CD205, DLL4, K14, MHCI and MHCII staining and merged channels with Hoechst. (F, G) WT mTOs cultured in the presence of RANKL showing K8, K14 and AIRE staining and merged channels with Hoechst. Right hand panel in G shows detail from boxed area of middle panel. (A-G) Each mTO shown is from a different well. n=3 independent biological replicates each with several technical replicates, where n is an independent biological replicate performed on a separate day, except for the MCHI, MHCII, β5t and the keratin staining where n=2. Hoechst reveals nuclei.

Live imaging of mTO showed that GFP, reporting *Foxn1*, was expressed in mTO throughout the 7 day culture period (the latest timepoint analyzed; Figure 4A, B), in contrast to 2-dimensional culture and submerged FTOC in which *Foxn1* is rapidly downregulated^56^. Counterstaining showed *Foxn1* was expressed in most TEC, while transcriptome analysis confirmed that *Foxn1* expression was maintained and thus that the persisting GFP signal reflected continued *Foxn1* transcription throughout the culture period (Figure 4A, Figure 5). Counting of cells segmented on nuclear staining (Figure 4C) revealed each mTO typically contained around 1000 cells.

**Figure 5:**
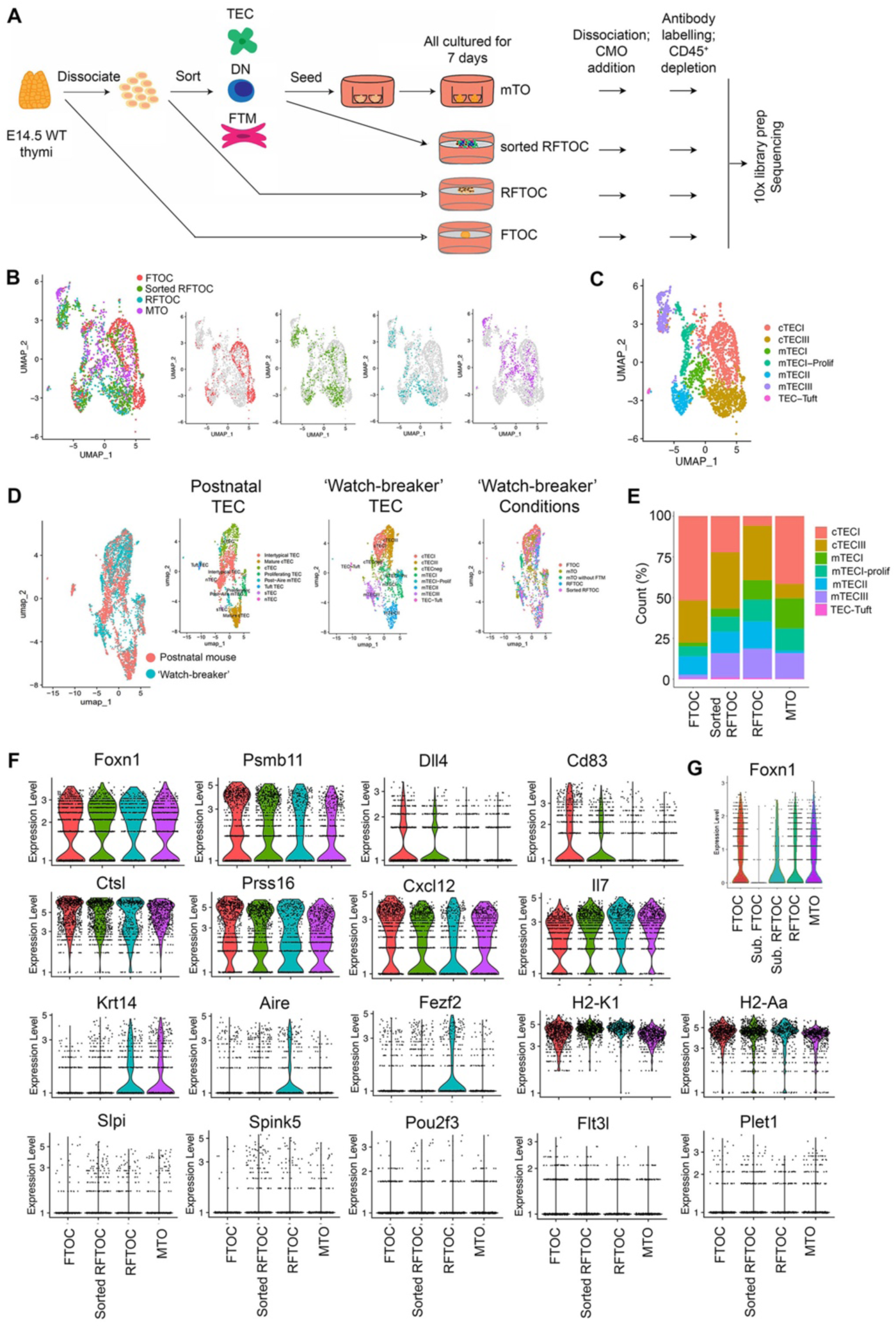
mTO contain all major TEC sub-types, determined by scRNAseq. (A) Schematic representation of experimental design for 10x scRNAseq data shown in (B-G). (B) UMAPs show combined data from the FTOC, RFTOC, sorted RFTOC and mTO conditions (left panel), plus the contribution of individual conditions to the combined dataset. (B-G) Only data for TEC are shown. (C) Unsupervised clustering revealing TEC subsets among combined dataset in (B). Annotation of clusters follows the naming convention in^57^. (D) UMAPs showing dimensionality reduction of the ‘Watch-breaker’ dataset combined with the postnatal TEC dataset from^59^. Combined data are shown in the left panel), and contribution of the Watch-breaker conditions and postnatal TEC to the combined dataset are shown in the middle and right panels respectively. (E) Graph shows proportional representation of clusters for each Watch-breaker condition, as shown. (F, G) Violin plots showing expression of a selected panel of genes, as shown, in the Watch-breaker conditions shown. See also Figures S6-13, See also Tables S3, S4 and S5.

Venus^+^ mTEC and dsRed^+^ cTEC present within the mTO clustered separately (Figure 4C, D), indicating that cTEC and mTEC regions had been established and were spatially segregated, while staining for cTEC and mTEC markers revealed DLL4, β5t, CD205, K8, K14, AIRE, MHCI and MHCII expression (Figure 4E,F). Note that some VENUS^+^ cells were found just adjacent to mTO (Figure 4C,D) and likely represent mesenchymal cells, based on cell and nucleus size and morphology. AIRE^+^ mTEC were present in ∼25% of mTO, with 1-2 AIRE^+^ cells typically detected per organoid by immunostaining after 7 days in culture; addition of RANKL to the culture medium increased both the percentage of mTO containing AIRE^+^ mTEC, and the number of AIRE^+^ mTEC per organoid by day 7 (Figure 4F).

FTM cells, identified by Vimentin staining, were present in all mTO (Figure 4A), where they preferentially associated with each other and exhibited two distinct distribution patterns: a single layer of cells separating two layers of TEC, or a central cluster of cells surrounded by TEC (Figure 4A) - with the first form predominating (11/14 versus 3/14 mTO scored).

Overall, these data show that mTO contain cTEC and mTEC expressing the expected sub-lineage restricted functional markers and are segregated rather than interspersed within the mTO structure. They further establish that mTO are suitable for live and fixed wholemount imaging applications, including timelapsing.

### Transcriptome analysis reveals that mTO TEC subpopulation complexity is comparable to that in RFTOC, FTOC and the native thymus

To further probe cellular complexity and phenotype, we performed single cell RNA sequencing analysis (scRNAseq). We set up nine ‘Watch-breaker’ conditions for analysis (Figure 5A, Figure S6): (i) FTOC, (ii) submerged FTOC, (iii) submerged RFTOC (unsorted cells), (iv) RFTOC (unsorted cells), (v) RFTOC without MEFs (unsorted cells), (vi) RFTOC in mTO proportions (sorted cells), (vii) unsorted mTO (i.e. mTO made without MEFs from unsorted E14.5 thymic cells), (viii) mTO+FTM (i.e. Hi FTM mTO), (ix) mTO-FTM (i.e. as for viii but without FTM). This allowed us to control independently for the effects of submerging, presence/absence of murine embryonic fibroblasts (MEFs), cell sorting, FTM, and varying input cell proportions. We were unable to recover any cells from condition (v) at the experimental endpoint (7 days).

Cells from all the remaining conditions were recovered and processed for flow cytometric analysis (CD45^+^ cells) or scRNAseq (CD45^-^ cells) as shown (Figure 5A; Figure S7; see Materials and Methods for details). Analysis of the CD45^+^ populations revealed poor thymocyte development in the sequenced mTO relative to the mean mTO performance, although all thymocyte populations analysed were present (Figures S8 and S9; see also Figure 2), indicating the sequenced mTO represented the lower end of the range of mTO outcomes. All other conditions performed as expected (Figures S8 and S9).

We sequenced the individual CD45-depleted cells from the eight conditions from which cells were recovered, using the 10x Chromium chip. Following initial quality control (Materials and Methods), the resulting dataset comprised 4919 single-cell transcriptomes, with a median of 4512 unique genes per cell. Unsupervised clustering across all cells that passed QC from all ‘Watch-breaker’ conditions, using the Seurat package in R, revealed a number of TEC clusters and a separate cluster of mesenchymal cells, as expected, along with a small cluster of thymocytes (Figure 5B; see Figure S10 for plots including mesenchymal cells and thymocytes, and all ‘Watch-breaker’ conditions). TEC clusters were identified as shown in Figure 5C and Figure S11, following the naming convention of Maehr and colleagues^57,58^: cluster ‘mTECI’ represents cells also referred to as ‘intertypical TEC’^59^, ‘jTEC”^60^ and ‘pre-Aire mTEC’^61^; clusters mTECII and mTECIII cells are often called ‘mature mTEC’ and ‘post-Aire mTEC’ respectively^59^. While cTEC are often split into three clusters - cTECI-III, with little difference between cTECI and cTECII^57^ - we found two cTEC clusters, which we called cTECI and cTECIII, characterised by low and high expression of MHC Class II (MHCII), respectively. The cTECIII cluster exhibited high expression of genes required in cTEC to support thymocyte development such as *Dll4* and *Psmb11*, as well as the ‘perinatal cTEC’ marker *Cd83*^59^. In human TEC, a cell type called ‘mcTEC’ or ‘PolyKRT TEC’, is marked by *Krt14* and *Krt15*^13,40^. While there is very little *Krt15* expression in the mouse thymus, *Krt14* expression is highest in mTECI cells (Figure 5C), suggesting mTECI is the mouse equivalent of mcTEC. The mTECI-proliferating cluster was very similar to mTECI but with high *Mki67* expression. Close correspondence between the cultured TEC and *ex vivo* TEC was demonstrated by clustering the ‘Watch-breaker’ datasets with those of Baran-Gale^59^; cells from the two datasets were interspersed in all TEC clusters (Figure 5D).

We first analyzed the FTOC, RFTOC, sorted RFTOC and mTO conditions, focussing on the TEC subpopulations in (Figure 5B,C,E-G). These conditions all contained cells in each of the TEC clusters identified, but differences in distribution across the conditions were observed including between FTOC and RFTOC (Figure 5B,C,E). mTO contained relatively few cTECIII and mTECII (although mTO cells were present in each of these clusters), but were well represented in all other clusters including mTECIII and TEC-Tuft and contained the highest proportion of mTECI- proliferating cells (Figure 5B-E). The low number of mTO cells in the mTECII (‘mature mTEC’) cluster was consistent with our immunohistochemical analysis and with the low numbers of DP thymocytes present in the sequenced mTOs (see Figure 5E and Figures S8 and S9 above; DP thymocytes provide RANKL required for mTECII development^62^). However, mTO mTECII cells clustered with mTECII from FTOC and RFTOC (Figure S12A, Table S5) indicating they are phenotypically normal.

We next analyzed expression of genes known to be important for TEC functionality, considering in our analyses differences in overall expression levels, and differences in expression levels in specific TEC subsets. All four conditions maintained *Foxn1* expression throughout the culture period, with the overall level of *Foxn1* in mTO at least equivalent to that in FTOC (Figure 5F, Table S4, 70% vs 72% *Foxn1*^+^ cells, adjusted p-value=1), consistent with reporter gene expression (Figure 4C). The four conditions also all expressed the cTEC markers *Dll4*, *Psmb11* (encoding β5t), *Prss16* (encoding TSSP), *Cxcl12*, and *Cd83* (Figure 5F), consistent with antibody staining (Figure 4E). None of these markers showed statistically significant expression in mTO across all other conditions, although some condition-to-condition comparisons were statistically significant (e.g. for *Psmb11* the adjusted p-value is <10^-6^ for mTO vs FTOC but 1 for mTO vs RFTOC, Table S4). This was also true for *Il7, Kitl* and *Plet1* and for the mTEC genes *Krt14*, *Fezf2*, *Spink5*, *Slpi* and *Pou2f3* (Figure 5F, Table S4). *Aire* was significantly downregulated in mTO versus all other conditions, however, this reflected the poor representation of mTECII as within that population (which expresses *Aire*); *Aire* expression was not significantly different between mTO and FTOC (Figure S12B; Table S5). In contrast, *H2-K1* and *H2-Aa* (encoding MHCI and II alleles) were also expressed in all conditions, but at slightly lower levels in mTO compared to each other condition (e.g. for *H2-K1* adjusted p-value<0.05 vs FTOC, 10^-53^ vs RFTOC and 10^-54^ vs sorted RFTOC, Figure 5F, Table S4); of note, this may reflect the low representation of mTECII in mTO since in mTECII the levels of *H2-K1* and *H2-Aa* were not significantly different for mTO compared to all other conditions (Table S5). Overall, compared to all other conditions put together the top downregulated genes in mTO were *B2m*, *H2-K1*, *Ifit1*, *Ubd*, *Bst2*, *H2-D1* and *H2-T23* (Table S4, mTOvsAll3), while in mTECII only *Stmn2* was differentially expressed across all conditions (Figure S12B, Table S5).

Expanding our analysis to submerged conditions revealed that submerged FTOC expressed extremely low levels of *Foxn1*, as expected^56^; and submerged RFTOC also expressed much lower *Foxn1* levels than RFTOC cultured at the air liquid interface (ALI) (Figure 5G). However as noted above *Foxn1* was not downregulated in mTO (also a submerged condition) relative to ALI FTOC or RFTOC (Figure 5G, see also Figure 4A). Differences in cellular composition were also observed between FTOC, RFTOC and their submerged counterparts (Figure S13A,B). Submerged FTOC and submerged RFTOC expressed lower levels of *Psmb11*, *Dll4*, *Aire*, *Fezf2*, *Slpi* and *Pou2f3* than their ALI counterparts (SFigure S12C), while as shown above (Figure 5F, Figure S13C) mTO expressed similar levels to ALI RFTOC and/or FTOC for all of these markers. Some differences were observed that might be attributable to cell dissociation. Most notably, *Dll4* and *Cd83* were downregulated and *Aire* and *Fezf2* upregulated in unsorted RFTOC versus FTOC that were otherwise cultured under identical conditions (Figure 5F). However, although consistent between unsorted RFTOC and mTO, these changes were not seen in sorted RFTOC. Examination of hypoxia induced genes (e.g. *Higd1a*) did not indicate a clear role for hypoxia in the differences observed (Figure S13D); indeed hypoxia-induced genes were expressed more highly in FTOC than in other conditions. No clear effects of cell sorting were evident (Figure 5F; note that in the sorted RFTOC condition the proportion of input cells is as for mTO, rather than as in unsorted RFTOC).

Collectively, these data show that mTO contain all major subtypes of TEC and that each subtype expresses the expected range of markers at ‘normal’ levels. Compared to FTOC and RFTOC, the most notable differences in mTO are the relative paucity of *Aire^+^*mTECII and slightly lower expression of MHC genes and interferon targets. Furthermore, our data reveal clear effects of submerging on gene expression in FTOC and RFTOC, but not in mTO.

### Transcriptome analysis reveals activation of interferon and antigen presentation pathways by FTM

To gain insight into the mechanisms regulated by FTM in TEC we also compared mTO with mTO minus FTM, revealing several key differences. Firstly, dimensionality reduction showed the vast majority of the ‘No-FTM’ mTO cells formed a distinct cluster adjacent to the cTECI region (Figure 6A), which we named ‘cTECneg’ due to the extremely low levels of both MHCI and MHCII (Figure 6A,B), with the remaining cells contributing to clusters mTECI, mTECI-Prolif and mTECIII (Figure 6A). None of the other conditions contributed to the cTEC-neg cluster. Secondly, the ‘No-FTM mTO’ condition expressed similar levels of *Foxn1* and *Il7* to mTO, FTOC and RFTOC and only slightly lower levels of *Psmb11*, *Cxcl12, Ctsl, Prss16, Plet1, Slpi* and *Spink5* than the other conditions. However, the ‘No-FTM mTO’ expressed markedly less MHCI (*H2-K1*), MHCII (*H2-Aa*), *Aire*, *Fezf2, Pouf2* and *Krt14* than mTO+FTM, and almost no *Dll4* or *Cd83* (Figure 6B,C, Tables S6 and S7). Notably, *Prss16*, *Dll4*, *Cd83*, *Tbata*, *Psmb11* and *Ccl25* are all among the high confidence FOXN1 target genes identified by ChIPseq^63^ and the differential expression of these genes in *Foxn1*^+^ cells in the ‘No-FTM mTO’ condition thus indicates that co-factors directly or indirectly regulated by FTM are required in addition to FOXN1 for their expression.

**Figure 6:**
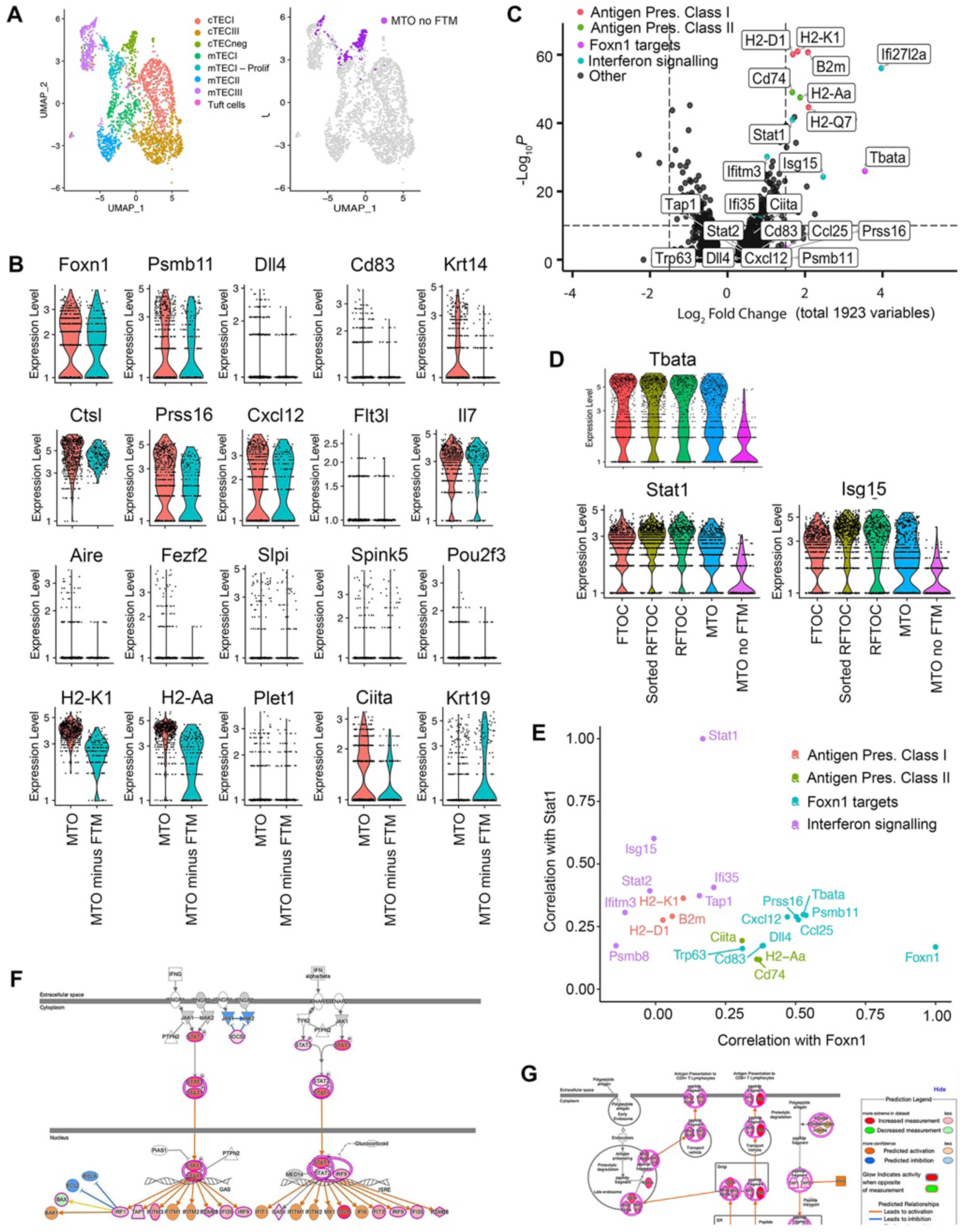
Effect of FTM on TEC populations and gene expression within mTO. (A) UMAPs show combined data from the FTOC, RFTOC, sorted RFTOC, mTO and mTO-FTM conditions (left panel), plus the contribution of the mTO-FTM condition to the combined dataset (right panel). (B-G) Only data for TEC are shown. (B) Violin plots showing expression of a selected panel of genes, as shown, in the Watch-breaker conditions shown. (C) Volcano plot showing differential gene expression between the mTO and mTO-FTM conditions; X axis, positive numbers are upregulated in mTO versus mTO-FTM. (D) Violin plots showing expression of selected IFN pathway genes, in the Watch-breaker conditions shown. (E) Plot shows correlation with *Foxn1* expression level and *Stat1* expression level of the differentially expressed genes highlighted in (C). (F, G) Images show Ingenuity pathway analysis of genes upregulated in mTO versus mTO-FTM, for the IFN signalling and antigen processing and presentation pathways. See also Figures S12-15, See also Tables S6 and S7.

The top differentially expressed genes between the mTO and No-FTM mTO conditions were *B2m*, *H2-D1*, *H2-K1, H2-Aa, H2-Q7* and *Cd74,* all members of the MHCI or MHCII antigen presentation pathways; the TEC regulator *Tbata;* and the interferon (IFN) stimulated genes (ISG) *Ifi27l2a* and *Isg15*, all of which were higher in the mTO condition (Figure 6C,D). Pathway analysis also highlighted the downregulation of the IFN signalling and Antigen Processing and Presentation pathways in the ‘No-FTM mTO’ versus mTO condition (Figure 6D-G). *Stat1*, a key regulator of IFN signalling implicated in regulation of antigen processing and presentation pathway components including *Ciita*, *Tap1, Tap2* and *Psmb8,* was among the most differentially expressed genes (Figure 6C-E). Targets of both type I and type II interferon signalling were also expressed more highly in the mTO with FTM condition, including *Irf1, Irf9, Tap1, Ifitm3* and *Ifi35.* as were additional antigen presentation pathway components including *Ciita* and *Tap1* (Figure 6C-G, Tables S6 and S7).

Examination of cells in the TEC and mesenchymal clusters of the FTOC, RFTOC and mTO conditions revealed almost no *Ifng* expression, some *Ifnz* expression in cTEC (in all conditions except mTO-FTM) and *Ifnb1* and *Ifnl2* expression in approximately 2% of mTECII, as reported elsewhere^64^ (Figure S14 and Table S5). This essentially rules out a direct effect of IFN signalling from FTM, leaving the possibilities that FTM is boosting IFN production in TEC themselves or in one or more intrathymic haematopoietic cell type. Of note is that MHC I expression in TEC has recently been shown to be independent of *Ifnar1* and *Ifngr1*, but dependent on *Ifnlr1, Aire, Stat1* and *Nlrc5*^65^, consistent with the hypothesis that interferon signalling is caused by type III interferons from mTECII.

Related, we observed two subpopulations within the mTECIII cluster, with all No-FTM mTO mTECIII cells belonging to one subcluster (Figure 6A). This subcluster expressed high levels of *Krt19, Notch1, Cldn3, Cldn4, Sox9,* and *Plet1*, markers of mTEC progenitors^66,67^, while both subclusters expressed the post-AIRE mTEC (mTECIII) markers *Slpi* and *Spink5*, which early mTEC progenitors do not. This raises the possibility that mTECIII contains an mTEC progenitor population as well as post-AIRE mTEC. In support of this hypothesis, the algorithm CytoTRACE2^68^ predicts the mTEC-progenitor-like subcluster of mTECIII to be less differentiated than the remaining subcluster (Figure S15).

## Discussion

Taken together, the data presented demonstrate that functional miniaturized thymic organoids can be generated from E14.5 fetal thymic cells in a microwell format suitable for medium throughput imaging and screening applications, including live imaging and time-lapsing. They establish the optimized cellular input requirements for these mTO and show they meet and surpass the minimal criteria for a functional thymic organoid. mTO can support T-cell lineage commitment and subsequent thymocyte differentiation to generate mature naïve SP4 and SP8 T-cells. They contain regions of cTEC and mTEC that express canonical sub-lineage specific markers (cTEC: DLL4^44,45^, CD205, CXCL12^46^, β5t^47^, Cathepsin L^48^; mTEC: RANK^69^, K14, AIRE^14^; TEC markers: MHCI and MHCII), Furthermore, global transcriptome analysis shows mTO contain all major cortical and medullary TEC subtypes, albeit with relative under-representation of mTECII compared to the early postnatal thymus – likely explained by the low numbers of DP thymocytes in the particular mTO sequenced. Through use of the mTO system to probe the role of FTM in thymic organoids, our data also identify a necessary, likely at least partly indirect, role for FTM in supporting differentiation and/or maintenance of most TEC subsets from E14.5 input cells and identify IFN signalling as a candidate mechanism for driving this crosstalk. These findings collectively raise several important issues, as discussed below.

Our findings introduce mTO as a new platform for exploration of thymus biology, compatible with both live imaging and medium throughput screening applications. In this system, each well of a 96 well plate contains at least 31 mTOs, enabling establishment of multiple replicates within each experimental condition while starting from a minimal number of input cells. Each mTO can be live-imaged individually, allowing parallel analysis of organoid, TEC and/or thymocyte behaviour over time in response to experimental manipulations including screening conditions.

mTO meet the formal definition of an organoid, namely “self-organized three-dimensional structures which mimic the key functional, structural and biological complexity of an organ .… derived from … stem cells or tissue-derived cells, including normal stem/progenitor cells …”^70^. They represent a step forward for analysis of thymic stromal biology and T-cell development, as the only currently available near-physiological models (FTOC and RFTOC) require large cell numbers for their establishment, precluding parallel testing of multiple conditions. In contrast, mTO are easier to establish, retain key physiological attributes of the fetal/perinatal thymus and can easily be used for live and wholemount imaging. As discussed below, our data already show the value of this platform for interrogating thymus biology (here, the role of FTM); beyond this, mTO are amenable to further development for genetic manipulation and modelling, and for high throughput screening protocols.

mTO can currently be produced in two formats – (i) based on dissociated but unsorted fetal TEC and (ii) based on dissociated and sorted fetal TEC reassembled using optimised proportions. The former is already amenable to medium throughput applications (e.g. to set up 96 wells each containing 31 unsorted mTO would require approximately fifteen timed matings), while the latter is currently suitable for smaller scale screening applications, e.g. enabling parallel testing of a smaller number of wells e.g. for image-based screening of multiple mTO within a single well per condition (in our hands, for ‘optimised mTO’, eight 96 wells can be established per ten timed matings). We expect this platform to develop rapidly, for instance by incorporating *in vitro* generated^37,38^ or expanded^39,40,42^ TEC.

In terms of ‘watch-breaking’ and ‘watch-making’, in the first part of our analysis we sought to determine which of the stromal cell types present in the native fetal thymus were required to generate functional mTO. Iterative testing revealed the requirement for both TEC and FTM, while omission of thymic EC did not affect the capacity of mTO to support thymopoiesis. From this, we conclude that in terms of stromal inputs the fetal thymus ‘watch’ cannot easily be broken: not only are all elements bar EC required to elaborate a functional mTO, the requisite stromal elements must be in physiological proportion (TEC:FTM was 2.5:1) for optimal function. The reason for this physiological ratio remains to be determined, but likely reflects the mechanisms through which FTM supports TEC and thymocytes^71-73^, including provision of extracellular matrix and of growth factors. The requirement for physiological proportionality however did not hold for thymocytes; optimal mTO outputs were achieved when the ratio of input TEC:DN cells was raised from 1:2 to 2:1, possibly reflecting reduction in niche availability due to the cell dissociation and reaggregation process. In support of this, RFTOC generated with optimised mTO cell input proportions supported thymocyte development better than FTOC and unsorted RFTOC in terms of output numbers for most thymocyte populations analyzed.

Our finding that FTM is an essential mTO component was expected. The absence of FTM-derived FGF7/10 leads to thymic hypoplasia^71^ and transplantation of FTM-depleted fetal thymic lobes leads to development of hypoplastic thymi^74^. Furthermore, FTM is an essential component of functional RFTOC, required to support early thymocyte development^72,73^. However, the effect observed herein was more severe than that in *in vivo* models lacking neural crest cell (NCC)-derived FTM. In *Pax3^-/-^* mice *(Pax3^Spl/Spl^* and *Pax3^Cre/Cre^*), which almost completely lack NCC-FTM, an ectopic and hyperplastic thymus develops until at least E14.5^75^, while transplantation of E9.0 NCC-free third pharyngeal pouch endoderm (3PPE) results in development of organised and functional thymi^76^ - consistent with the ability of transplanted mesenchyme-stripped thymic lobes to co-opt mesenchyme from the transplant site^77^. Our current findings indicate the near complete failure of mTO generated without FTM to support TEC development and/or maintenance, and the complete failure to support thymocyte development. This was surprising, since the DN input cells for mTOs included CD44^-^CD25^+^ (DN3) and CD44^-^CD25^-^ (DN4) cells, while the requirement for FTM in RFTOC has been mapped to the CD44^+^CD25^+^ (DN2) stage of thymocyte development with later stages being FTM-independent^72^. Furthermore, it did not result simply from failed reaggregation (see Figure S7).

Examination of the gene expression profile of mTO-FTM cells revealed equivalent levels of *Foxn1* to mTO+FTM cells (and all other conditions), ruling out loss of FOXN1 as the underpinning cause of its lack of functionality. Notably, despite this, the levels of some known direct transcriptional targets of FOXN1^63^, including *Dll4, Cd83* and to a lesser extent *Cxcl12*, *Psmb11, Ctsl* and *Prss16* were downregulated in mTO-FTM cells. While this may reflect the absence of cell types that normally express these markers, it further suggests that co-factors in addition to FOXN1 are required for normal expression of these genes and that these are limiting in cTEC-neg and mTECIII cells, and/or that accessibility of these loci is regulated in a TEC sub-type specific manner. This observation is consistent with our previous finding that not all direct FOXN1 targets are upregulated in every TEC upon overexpression of functional FOXN1 *in vivo*^78^, and with the specific expression of *Dll4*^44^ and *Psmb11*^79^ in cTEC.

Analysis of gene expression in the mTO+FTM versus mTO-FTM conditions revealed the most significant difference to be a marked downregulation of Type I IFN-regulated genes, including *Stat1*, MHCI (*H2-K1*), MHCII (*H2-Aa*) and multiple members of the Class I and Class II antigen processing and presentation pathways, suggesting that lack of FTM-regulated IFN signalling might explain the failure of mTO-FTM. However, while receptors for all interferons (*Ifnar1*, *Ifnar2*, *Ifngr1, Ifngr2* and *Ifnlr1)* were broadly expressed across all ‘Watch-breaker’ conditions, there were essentially no interferons observed in FTM and the only interferons observed in TEC were *Ifnz* (which was expressed by a few cTEC in FTOC, RFTOC and mTO) and *Ifnb1*, *Ifnl2* and *Ifnl3* (which were expressed in ∼2% of mTECII in all conditions with mTECII, as expected^64^).

IFN expression in the thymus is thought to be restricted to AIRE^+^ mTEC (mTECII)^64,65^ with *Aire, Ifnlr1, Stat1* and *Nlrc5* dependent expression of type III interferons regulating MHCI expression^65^. Although IFNα expression has been reported in human thymic plasmacytoid dendritic cells (pDC)^80^, as well as IFNγ expression in thymic iNKT-cells and eosinophils^81^, the expression of MHC I in the TEC was only dependent on the type III interferon receptor *Ifnl1* and not *Ifnar1* or *Ifngr1*^65^. Similarly, *Ifnlr1* was shown to play a bigger role than *Ifnar1* or *Ifngr1* in the maturation of thymic DC1s, macrophages, B cells and Treg cells^64^. In the mTO without FTM, the IFN-regulated gene signature was strongly downregulated in all cell types. This suggests IFN signalling may play an earlier, previously undefined role, in TEC development. Alternatively, it may simply reflect the absence of mTECII, as this cell-type (and some cTEC in our data) is normally responsible for IFN production. Since FTM has been shown to regulate proliferation of fetal TEC^74^ and differentiation of stem/progenitor cells is proliferation-dependent in at least some other lineages^82,83^ it is also possible that the absence of FTM leads to a lack of TEC proliferation, resulting in the absence of more differentiated TEC phenotypes. However, inspection of cell cycle regulators did not reveal an obvious deficit in the mTO-FTM cells (not shown). Thus, on balance, we favour a role for IFN signalling pathway in TEC differentiation and/or maintenance.

The inability of the ‘No-FTM’ mTO to support thymopoiesis could in part be explained by the very low levels of *Dll4* expression in this condition. This was puzzling, as *Dll4* is a known direct transcriptional target of FOXN1^63^, yet *Foxn1* levels were near-normal in mTO-FTM. VEGF is known to act via FOXC1 to regulate *Dll4* expression in endothelial cells^84^. *Foxc1* was expressed in some TEC across all conditions analyzed, thus the observed downregulation of *Vegf* in the mTO-FTM condition provides a possible explanation for the loss of *Dll4*.

Thymic organoids have enormous therapeutic potential, both for generating positively and/or negatively selected T-cells and acting as model systems for understanding thymic involution and regeneration. Three-dimensional cell-line based cultures, called Artificial Thymic Organoids (ATO) have already been used to generate CAR T-cells^85^ and mature T-cells from bone-marrow organoid derived hematopoietic cells^86^. As ATO lack TEC-specific functions required for beta, positive selection and tolerance induction, deployment of bona-fide thymic organoids such as mTO should substantially improve these systems. The establishment of a medium/high-throughput thymic organoid system, described here, will also be useful for mechanism interrogation, which will aid in the development of serum and feeder free thymic organoids, as well as treatments aimed at regenerating thymic tissue. Furthermore, thymic organoids have the potential to be patient-specific, e.g. if derived from patient PSCs, allowing for patient-specific TCR repertoire selection: we expect the mTO system to be readily adaptable to human fetal thymus and PSC-derived TEC inputs.

## Disclosure of Potential Conflicts of Interest

The authors declare that the research was conducted in the absence of any commercial or financial relationships that could be construed as a potential conflict of interest.

## Author contributions

**VM, PR, THe, SP:** Conception and design, collection and assembly of data, data analysis and interpretation, manuscript writing. **THu, JM:** Conception and design, collection and assembly of data. **JS, AC, JC-W, QG, YW:** Collection and assembly of data. **JC, ML:** Conception and design. **AB:** Provision of study materials. **GA:** Conception and design, Provision of study materials, data analysis and interpretation. **CCB:** Conception and design, financial support, provision of study material, data analysis and interpretation, manuscript writing, final approval of manuscript.

## Data Sharing Statement

The datasets generated for this study are available on request. The codes for the analyses shown in Figs 5 and 6 and related Supplemental data are available at: https://github.com/samIndeed/Watchbreaker.

## Funding

The research leading to these results received funding from the School of Biological Sciences, University of Edinburgh (TH, AC); the UKRI-Biotechnology and Biological Sciences Research Council through an industrial CASE studentship (award number BB/MO16412/1; CCB, PR) and EASTBIO DTP studentship (QG), and through a UKRI-BBSRC/NC3Rs grant (award number NC/X002470/1; CCB, VM); the Darwin Trust (JM); the European Union Seventh Framework Programme (FP7/2007-2013) collaborative project ThymiStem under grant agreement number 602587 (PR, TH, CCB); the UKRI-Medical Research Council through a DTP in Precision Medicine studentship (award number MR/N013166/1) (JS); and the Wellcome Trust collaborative award 211944/Z/18/Z (VM, PR, TH, YW, SP, THu, JC, GA, ML, CCB). GA is supported by a UKRI-MRC Programme Grant (MR/T029765/1).

## Acknowledgements

We thank C. Cryer and F. Rossi (IRR, University of Edinburgh) for cell sorting, M. Vermeren (IRR, University of Edinburgh) for imaging, and the BVS staff for animal care.

## Supplemental Information

**Document S1. Figures S1–S15 and Tables S1-3**

**Table S4-7**. Excel file containing additional data too large to fit in a PDF, related to Figures 5 and 6.

## Materials and Methods

### Mice

Foxn1^GFP^ ^52^, Rank-Venus^53^ and Cxcl12^dsRed^ ^55^ mice were as described. C57BL/6 mice were used for isolation of fetal thymic cells except where otherwise specified. For timed matings, noon of the day of the vaginal plug was taken as day 0.5 of embryonic development (E0.5). All animals were housed and bred in University of Edinburgh animal facilities. All experimental procedures were conducted in compliance with the Home Office Animals (Scientific Procedures) Act 1986, under project licence PEEC9E359 to V. Wilson. Primers used for genotyping were as shown in Table S1. All controls were littermates unless otherwise stated.

### Thymus dissociation

Microdissected fetal thymi were dissociated for 5 minutes in TrypLE Express Enzyme (Life Technologies 12604013) in an Eppendorf Thermomixer (1400 rpm, 37°C) followed by trituration with a 25G syringe ten times. Cell suspensions at 4°C were washed in 2% FCS FACS buffer, resuspended as required and filtered through a 70μm cell strainer (Corning) to remove clumps.

### Tissue culture

mTO, RTOC and FTOC were cultured in Advanced DMEM/F12 (Life Technologies, 12634010), 2% FCS (Life Technologies), 1% penicillin (10,000 units/ml)/streptomycin (10,000 μg/ml) (Invitrogen, 15140-122), 1% GlutaMax supplement (Life Technologies, 35050061), 1% non-essential amino acids (NEAA) (Invitrogen, 11140-036) (referred as 2% media herein). All cell manipulations were performed in a laminar flow sterile hood using sterile technique. Cell culture plastic ware was supplied by Iwaki. All solutions were tested for sterility and warmed to 37°C prior to use. Cells were examined using an inverted microscope (Olympus CK2).

### Generation of mTO

The required cell populations were obtained by dissociation of E14.5 fetal thymi followed by flow cytometric cell sorting if required (TEC, DAPI^-^TER119^-^CD45^-^EpCAM^+^; double negative thymocytes [DNs], DAPI^-^Lin^-^CD45^+^EpCAM^-^; FTM, DAPI^-^TER119^-^CD45^-^EpCAM^-^ PDGFRαβ^+^; endothelial cells [ECs], DAPI^-^TER119^-^CD45^-^EpCAM^-^PDGFRαβ^-^CD31^+^; see Figure S1), pelleted, resuspended in 2% media and counted using the BioRad Cell Counter with the addition of Trypan Blue. Based on the cell counts, an appropriate volume from each population were pipetted into each well one-by-one or after prior mixing. Separate suspensions were created for each experimental condition. A maximum of 40μl per 96 well was gently dropped into the Gri3D^®^ 96well plates (SunBioscience, Gri3D-96-S-8P) and left to reaggregate for 15 minutes, after which the wells were filled with medium to a total volume of 200μl. Medium was refreshed every other day using the reservoir well attached to each well. mTOs were cultured from 7 to 14 days depending on the experiment.

### FTOC and RFTOC

were generated as previously described^87^. The compacted cell pellets were extruded at the liquid-gas interface onto a polycarbonate filter paper raft floating on 1ml of medium in 24 well plates and cultured for 7 days. Medium was refreshed every other day. For FTOC, each whole intact lobe was placed on a filter membrane with the help of a microscope and cultured for 7 days as above.

### Flow Cytometry

Cells were processed for flow cytometric sorting and analysis as previously described^34,49^. Compensation controls were performed using beads. Sorting and analysis gates were set using FMOs. Sorting was performed using a BD FACS Aria II running FACS Diva 4.1 (BD Biosciences). FACS analyses were performed on a FACSCalibur (BD Bioscience) or Novocyte running NovoExpress 1.3.0 (ACEA) at the CRM, University of Edinburgh. All post-acquisition analysis was performed with FCSexpress 7 (De Novo Software) software.

### Immunohistochemistry

All fixation and staining steps were carried out in the imaging bottom Gri3D^®^ plates (Sun Bioscience, Gri3D-96IBI-S-8-800) such that the mTOs were kept intact. mTOs were washed (3x10 minutes) with PBS to remove excess medium, fixed in 4% PFA (Fisher Scientific UK Ltd, 15670799) for 20 minutes at room temperature (RT) while shaking slowly, and washed with 0.1%Triton (VWR, X100-100ML) in PBS/0.2% Sodium Azide (PBST). For the preparation of sections, mTOs were stained for HOECHST (1:1000) for 15 minutes at room temperature. mTOs were then embedded in 2% agarose, by incubating for 5 minutes at 65°C with the concentration gradually increasing from 0.125% to 2% in five incubation steps, then left to solidify and scooped out of the 96well in one piece. Further submersion in 2% agarose allowed preparation of 200μm Vibratome sections (Leica VT1000 S Vibrating blade microtome). mTOs (or sections) were then blocked in 10% goat serum in PBST for 4-6 hours at RT, incubated in primary antibody solution (optimized concentration in 1% goat serum in PBST) for up to 48h at RT, washed in PBST (3x15 minutes), incubated in secondary antibodies (optimized concentrations in 1% goat serum in PBS only) and HOECHST (1:1000) for 8-24h at RT, then washed in PBS (3x15 minutes). In the case of a dim HOECHST signal, another 1:1000 dilution was applied. 70μl RapiClear 1.49 (www.sunjinlab.com, RC149001) was then added to each well at least 15 minutes before imaging. For live imaging, mTOs were kept in the GRI3D^®^ plates and imaged using the Opera Phenix^®^ Plus High-Content Screening System in 5% CO_2_ at 37°C. Fixed and stained mTOs were imaged similarly.

### Antibodies

The antibodies used for immunohistochemistry and flow cytometry were as listed in Table S2.

### Image analysis

All post-acquisition analysis was performed using Image Artist software or ImageJ v1.53c. For the segmented image in Figure 4C, the images presented were blind deconvolved using the Richardson-Lucy algorithm. Initial PSF guess was computed using Gibson– Lanni model (https://github.com/MicroscPSF/MicroscPSF-Py)^88^.

Cells were segmented using an automated algorithm based on U-Net neural network architecture^89^. This network predicted two channels, namely binary mask of cells and distance to the cell edge. The Watershed algorithm was used to segment the binary mask into individual cells using predicted distance as energy^90^. The images fed to the neural network were not deconvoluted and were normalized to the range [0,1]. Deconvolved images were used for measurement of mean intensities for individual cells. Values were expressed as average values withing a segmented mask of each cell.

### Single cell RNA-seq

*Processing of samples for library preparation, library preparation and sequencing:* Cells from each condition were resuspended in 90μl FACS buffer (PBS without Ca^2+^ and Mg^2+^, with 2% FCS), 0.45μl FC block was then added and incubated for 10 minutes on ice. Each condition was incubated with barcode tagged antibodies against MHCII, CD40, CD80 and EpCAM, with biotinylated UEA1, and with α-CD45-magnetic beads (Table S3) for 15 minutes on ice, then washed once with 1ml FACS buffer and centrifuged at 500rcf for 5minutes. Each condition was hashed with 100μl lipid anchored Cell Multiplexing Oligos for sample multiplexing (CMOs, 10x Genomics)(see Figure S6). Samples were then passed through LS Columns to deplete CD45^+^ cells according to the manufacturer’s instructions. Cells eluted from the column were spun down and resuspended in 50μl FACS buffer, of which 10μl was taken for cell counting. Approximately 1-5000 cells were pooled from each condition, and were then filtered and loaded on a Chip G (10X Genomics). Cell-bead encapsulation was performed using the Chromium X (10x Genomics). Gene expression and multiplexing libraries were prepared following the manufacturer’s instructions using the Chromium Next GEM single cell reagent kit 3’ V3.1 (dual index) with Feature Barcoding technology for Cell Multiplexing (CG000388, 10x Genomics). Libraries were quantified using a Bioanalyzer DNA High-sensitivity Kit (Agilent Technologies), pooled at a molar ratio of 10:1 between gene expression and multiplexing libraries, and sequenced on an Illumina NextSeq 2000 P3 flow cell (100 cycles configuration) to obtain approximately 1.4B paired-end reads.

*QC and analysis:* FASTQ files were processed using Cell Ranger (10x Genomics) and count matrices (for RNAseq, CITEseq and CMO multiplexing) were analyzed using the R package Seurat. Deconvolution of the CMO tags revealed multiplets (318), where tags were shared between cells. These, plus unassigned (360) and blank (122) reads, were excluded from further analysis. The remaining captured cells were filtered such that only cells with >2000 RNA features were retained for downstream analysis. Normalisation was carried out using the Seurat function SCTransform with regression on mitochondrial gene transcript percentages. After initial clustering, a cluster with high levels of stress markers (*Ddit3* and *Herpud1*) was identified and excluded from further analysis. GO enrichment analysis was performed in R with the enrichR package, using the “GO_Biological_Process_2021” database. The Seurat function *DEenrichRPlot* was used with “max.genes=1000”. DE gene analysis was carried out in R using the Seurat function *FindMarkers* with default settings.

### Statistics and experimental design

For all scRNAseq analyses, n represents the number of independent biological experiments. For all other experiments, n represents an independent biological replicate. At least three biologically independent replicates were performed for each condition with the exception of the antibody staining where n=1 in some cases. No statistical method was used to predetermine sample size, the experiments were not randomized, and the investigators were not blinded to allocation during experiments and outcome assessment. There were no limitations to repeatability of the experiments. No samples were excluded from the analysis.

## Data availability

The datasets generated for this study are available on request. The codes for the analyses shown in Figures 5 and 6 and related Supplemental Information are available at: https://github.com/samIndeed/Watchbreaker.

## Notes

### Competing Interest Statement

The authors have declared no competing interest.

