## Supplemental Figures and Tables for "Watch-breaker: establishment of a microwell array-based miniaturized thymic organoid model suitable for high throughput applications"

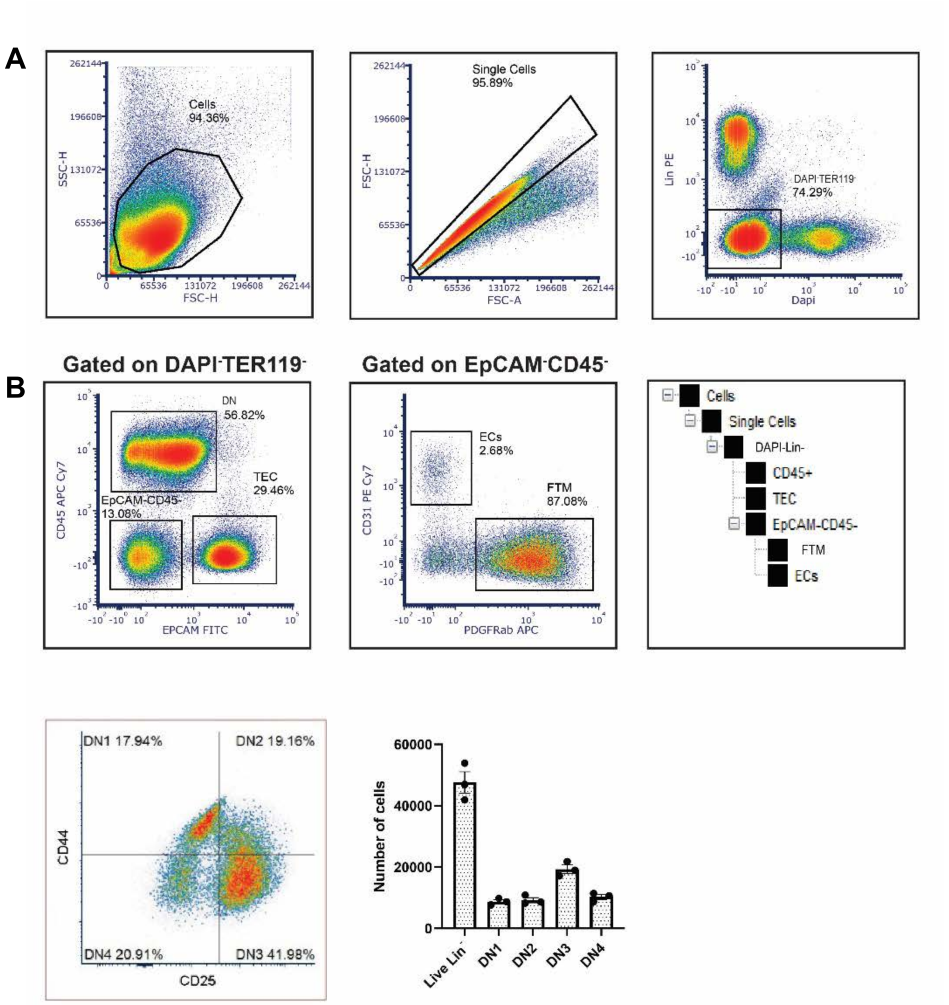


| | Live Lin$^-$ | DN1 | DN2 | DN3 | DN4 |
|---|---|---|---|---|---|
| Embryo1 | 41906 | 7736 | 8264 | 17192 | 8714 |
| Embryo2 | 53917 | 9817 | 10889 | 21804 | 11407 |
| Embryo3 | 46998 | 8695 | 8636 | 18998 | 10669 |
| Mean | 89.5 ± 0.04% | 18.39 ±0.09% | 19.43 ± 0.54% | 40.63 ±0.20% | 21.55 ±0.58% |

**Figure S1. Cellular composition of the E14.5 thymus.** A) Plots show gating strategy for isolating TEC, DN thymocytes, FTM and ECs from E14.5 thymic lobes. TEC were sorted from single cells as DAPI$^-$TER119$^-$CD45$^-$EpCAM$^+$ cells, DNs as DAPI$^-$Lin$^-$CD45$^+$EpCAM$^-$ cell, FTM as DAPI$^-$TER119$^-$CD45$^-$EpCAM$^-$PDGFRαβ$^+$ cells and ECs as DAPI$^-$TER119$^-$CD45$^-$EpCAM$^-$PDGFRαβ$^-$CD31$^+$ cells. B) Representative plot showing the CD44 versus CD25 subset profile, absolute cell numbers and percentages for E14.5 DAPI$^-$Lin$^-$CD45$^+$ cells. Lin = $\alpha$-CD4, $\alpha$-CD8, $\alpha$-CD11b, $\alpha$-CD11c, Gr1, NK1.1, B220, TER-119, $\alpha$-EpCAM.

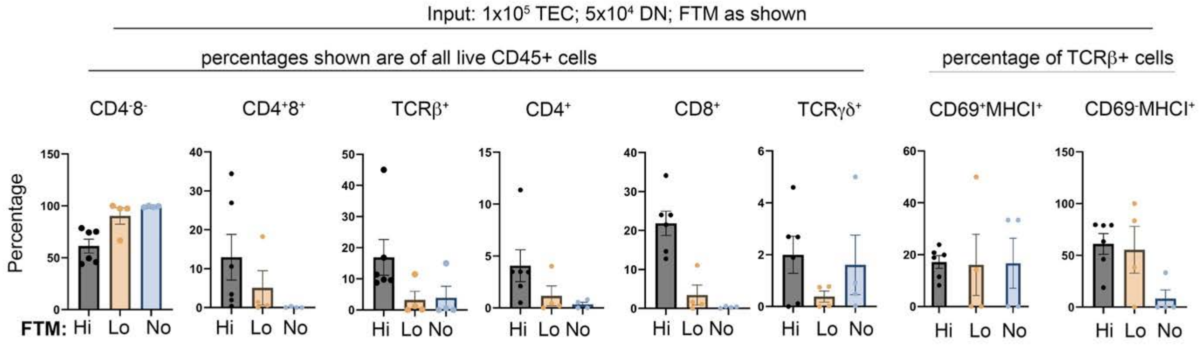


**Figure S2. Effect of FTM on mTO outputs.** mTO were established from E14.5 thymi using the following input conditions: Hi FTM (1x$10^5$ TEC, 5x$10^4$ DNs, 4x$10^4$ FTM), Lo FTM (1x$10^5$ TEC, 5x$10^4$ DNs, 1.5x$10^4$ FTM) and No FTM (1x$10^5$ TEC, 5x$10^4$ DNs, 0 FTM ) and were cultured for 14 days before analysis of thymocyte populations shown. Plots show mean±SEM of percentage for each subset. Each data point represents the cells harvested from one microwell. Percentages shown are after gating on the populations indicated. N = at least 4 independent biological replicates. DN ($CD4^-CD8^-$), DP ($CD4^+CD8^+$), ($CD3ε^+TCRβ^+$), SP4 ($CD4^+$), SP8 ($CD8^+$), $CD69^+MHCI^+$, $CD69^+MHCI^-$, γδ T cells ($CD3ε^+TCRγδ^+$).

A)

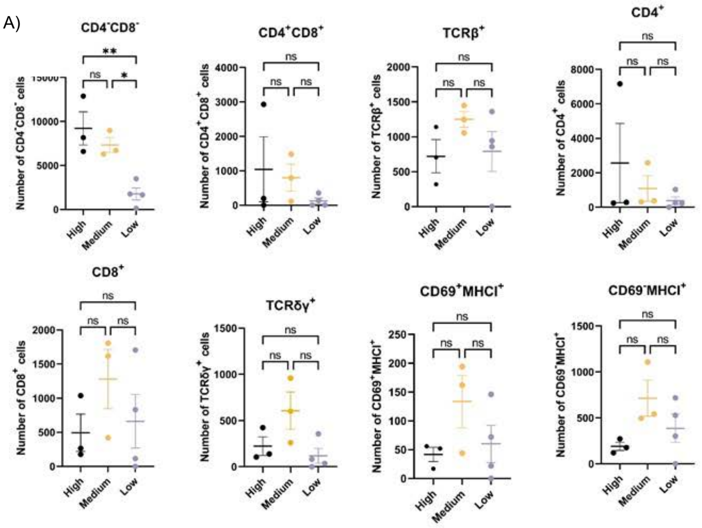


B)

| | Single cells | Live CD45$^+$Lin$^-$ | TCRβ$^+$ | CD69$^+$ MHCI$^+$ | CD69$^-$ MHCI$^+$ | TCRγδ$^+$ | SP4 (CD4$^+$) | DP | SP8 (CD8$^+$) | DN |
|---|---|---|---|---|---|---|---|---|---|---|
| **High number** | 73.7± 27% | 76.4± 4.9% | 5.3± 1% | 5.9± 1% | 32.2± 11.8% | 1.6± 0.5% | 17± 14.3% | 6.1± 5.5% | 3.4± 1.35% | 73.6± 12% |
| **Medium number** | 78.1± 5.2% | 77.5± 3.15 | 12± 1.4% | 10± 3.3% | 54.2± 9% | 5.8± 2.1% | 11.3± 8.1% | 7.1± 3.1% | 12.1± 4.1% | 70± 1.6% |
| **Low number** | 73.8± 14.4% | 68.2± 13.5% | 21.6± 7.5% | 5.4± 2.5% | 36± 12.9% | 2.5± 1.3% | 12.2± 8.4% | 2.8± 1.6% | 15.4± 7.8% | 69.71± 10% |

**Figure S3. Effect of total cell number on mTO outputs.** mTO were established using the same ratio but different absolute numbers of input cells per well (1.5, 1 or 0.5 x the Hi FTM condition numbers) and were cultured for 14 days before analysis of thymocyte development for the subsets shown. A) Graphs show mean± SEM of absolute cell numbers recovered per well for the input populations. N=3 independent biological replicates, one with two technical replicates for the Low proportion condition. B) Table shows cell mean±SEM of percentages of parent gates of defined thymocyte subsets present in High, Medium, Low input number mTOs. All data shown cell numbers after gating on live CD45$^+$ cells except the CD69$^+$MHCI$^+$ and CD69$^-$MHCI$^+$ populations which are gated on TCRβ$^+$ cells. Statistical analysis was one-way ANOVA or Kruskal-Wallis rank test based on Shapiro-Wilk normality test. ns not significant; $p > 0.05$. DN (CD4$^-$CD8$^-$), DP (CD4$^+$CD8$^+$), (CD3ε$^+$TCRβ$^+$), SP4 (CD4$^+$), SP8 (CD8$^+$), CD69$^+$MHCI$^+$, CD69$^+$MHCI$^-$, γδ T cells (CD3ε$^+$TCRγδ$^+$).

A)

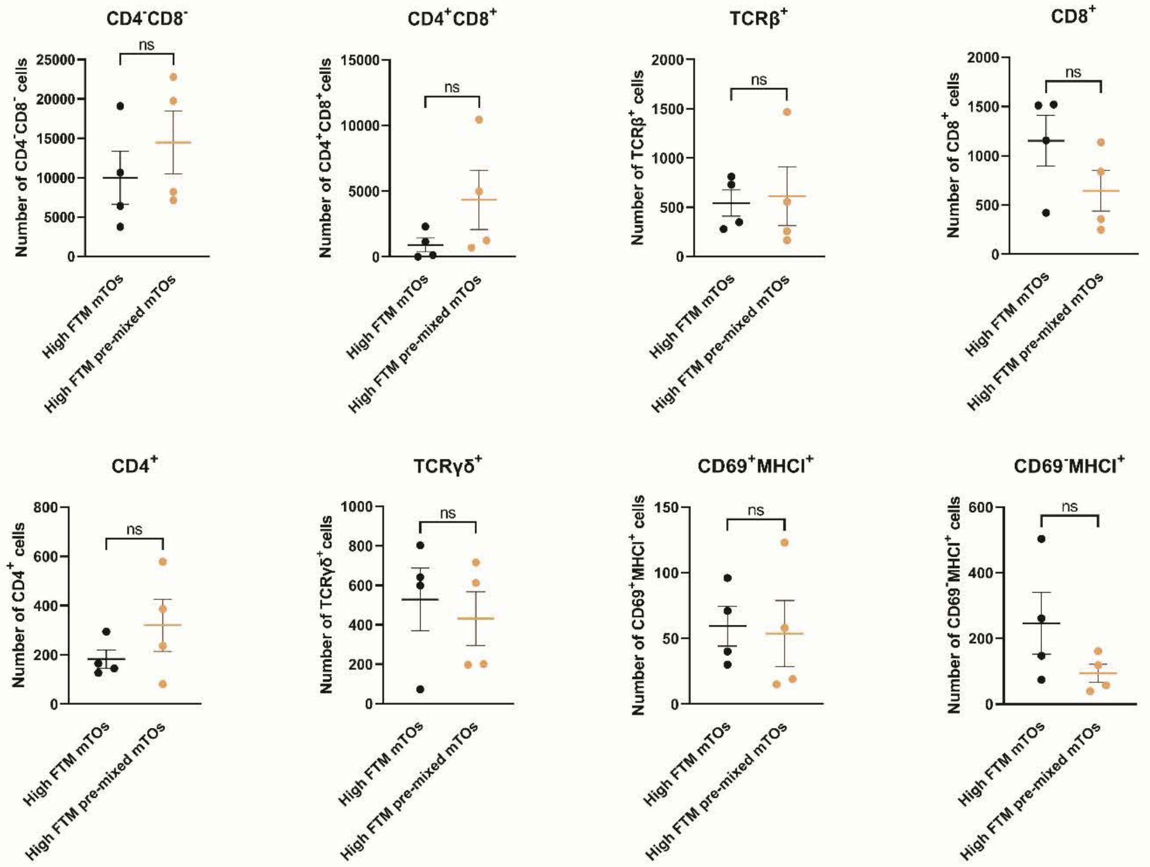


B)

| | Single cells | Live CD45+Lin- | TCRβ+ | CD69+ MHCI+ | CD69- MHCI+ | TCRγδ+ | SP4 (CD4+) | DP | SP8 (CD8+) | DN |
|---|---|---|---|---|---|---|---|---|---|---|
| High FTM | 81.4± 6.3% | 74.3± 2% | 5.3± 1.7% | 11± 0.5% | 41.6± 7.5% | 4.5± 1.5% | 1.7± 0.3% | 8± 4.8% | 10.6± 2.8% | 79.7± 4% |
| High FTM pre-mixed | 82.6± 7.5% | 78.9± 4.5% | 2.7± 0.5% | 8.8± 0.6% | 19.4± 2.9% | 2.2± 0.12% | 1.6± 0.28% | 17.5± 4.7% | 3.3± 0.1% | 77.6± 4.5% |

**Figure S4. Effect of pre-mixing input cells on mTO outputs.** mTO were established from E14.5 thymi using High FTM conditions with or without pre-mixing of input populations (in the without pre-mixing condition the input cell types were added sequentially) as indicated and were cultured for 14 days before analysis of thymocyte development for the subsets shown. (A) Graphs show mean± SEM. High FTM and High FTM pre-mixed had the same cellular composition ($1x10^5$ TEC, $5x10^4$ DNs, $4x10^4$ FTM). Each data point represents the cells harvested from one microwell. (B) Table shows mean±SEM for percentages of parent gates of the thymocyte subsets shown in High FTM and High FTM pre-mixed mTOs. N=4 independent biological replicates. Statistical analysis was by unpaired t-test or Mann- Whitney rank test based on Shapiro-Wilk normality test. ns, not significant; $p > 0.05$. DN ($CD4^-CD8^-$), DP ($CD4^+CD8^+$), ($CD3ε^+TCRβ^+$), SP4 ($CD4^+$), SP8 ($CD8^+$), $CD69^+MHCI^+$, $CD69^+MHCI^-$, γδ T cells ($CD3ε^+TCRγδ^+$).

A)

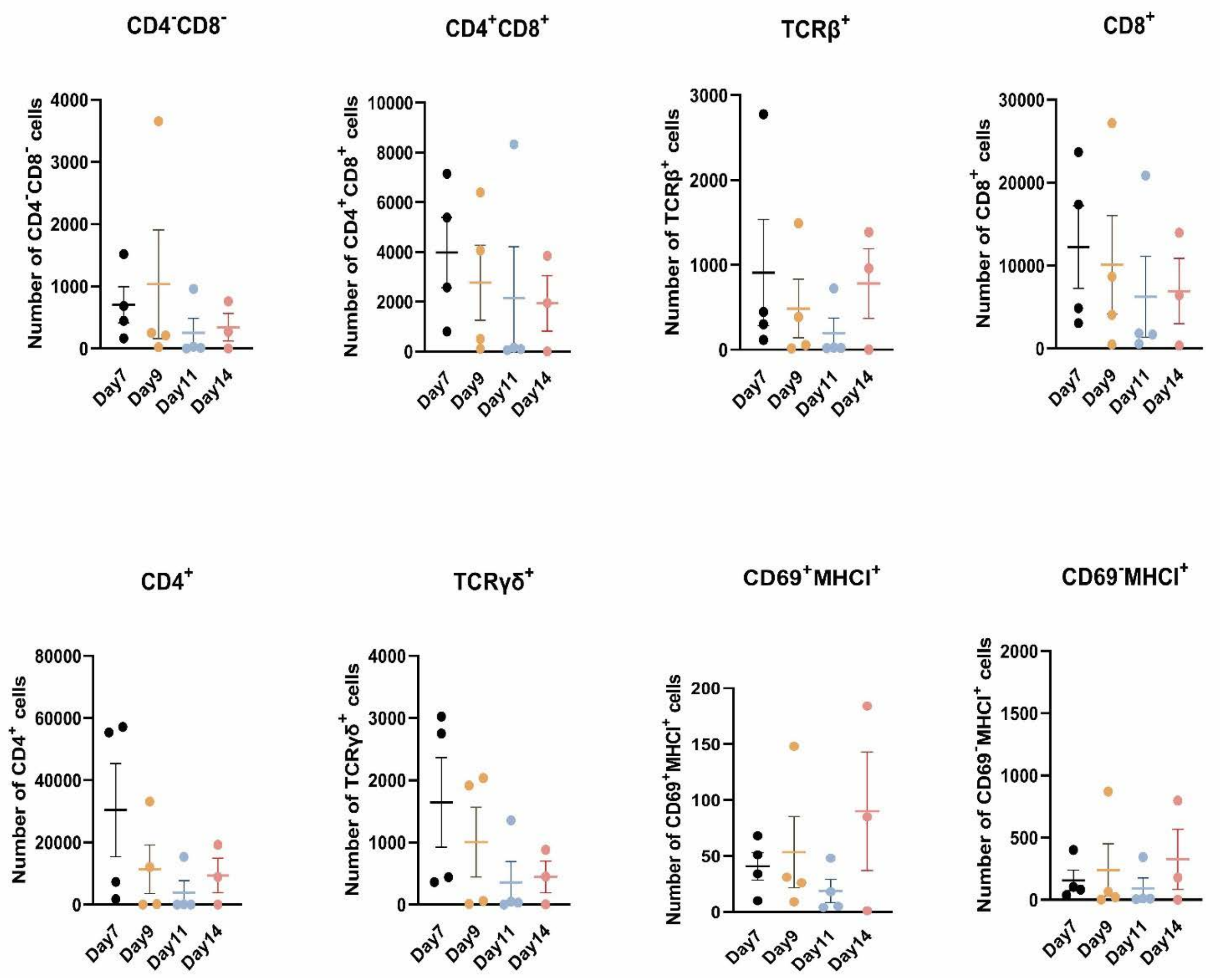


**Figure S5. Effect of culture period on mTO outputs.** mTO were established from E14.5 thymi using High FTM conditions and cultured for the time-periods indicated before analysis of thymocyte development for the subsets shown. (A) Graphs show mean± SEM. Each data point represents the cells harvested from one microwell. N=4 independent biological replicates. DN ($CD4^-CD8^-$), DP ($CD4^+CD8^+$), ($CD3\varepsilon^+TCR\beta^+$), SP4 ($CD4^+$), SP8 ($CD8^+$), $CD69^+MHCI^+$, $CD69^+MHCI^-$, γδ T cells ($CD3\varepsilon^+TCR\gamma\delta^+$).

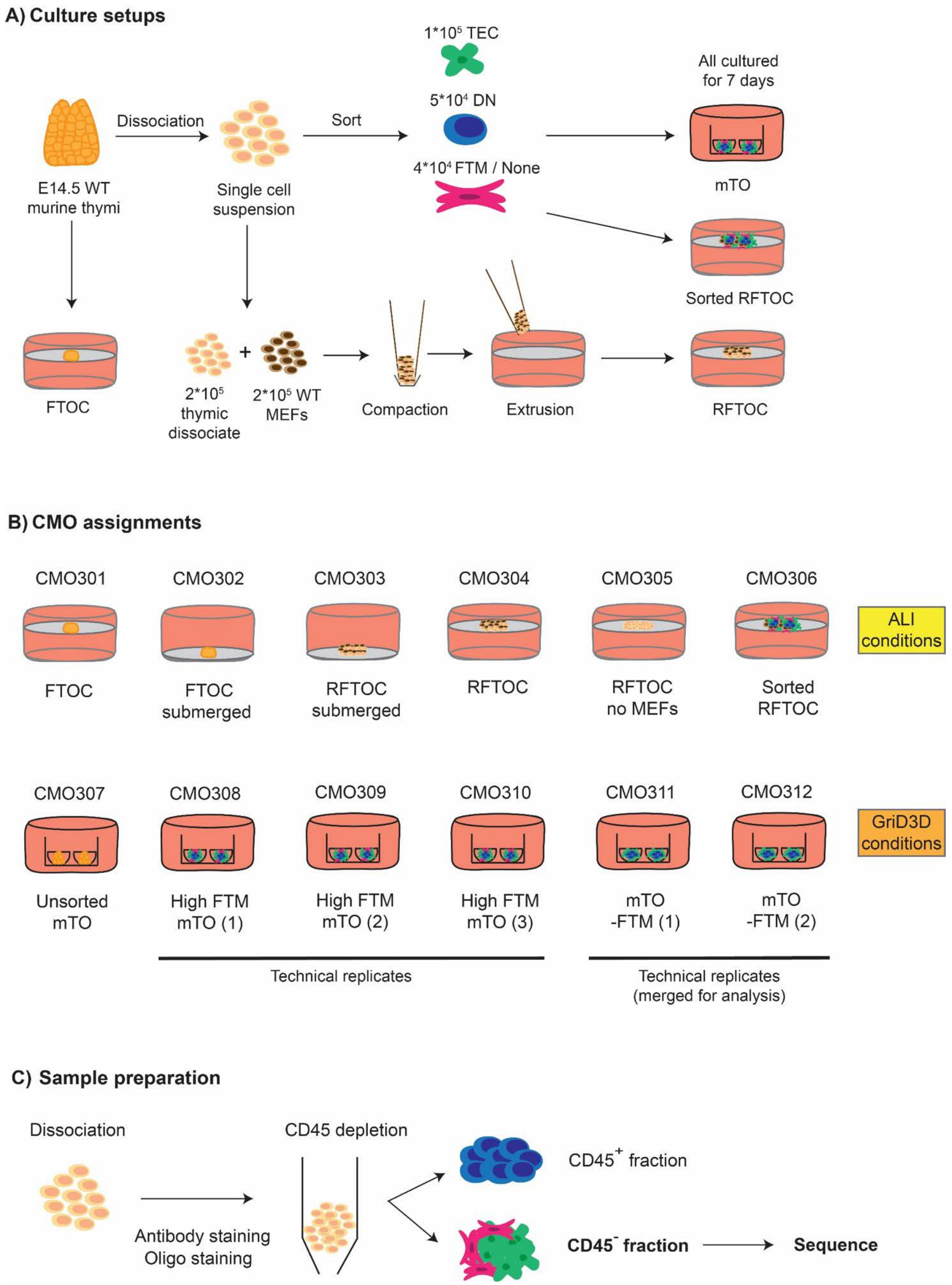


**Figure S6. Experimental design for scRNAseq analysis.**

A)

| | Condition | Sort | FTM | MEFs | Plate | Total cell input | Submerge | TEC | FTM | DNs |
|---|---|---|---|---|---|---|---|---|---|---|
| CMO301 | FTOC | No | Yes | No | ALI | 1 lobe = $2*10^5$ | No | | | |
| CMO302 | FTOC submerged | No | Yes | No | ALI | 1 lobe = $2*10^5$ | Yes | | | |
| CMO303 | RFTOC submerged | No | Yes | Yes | ALI | $4*10^5$ | Yes | | | |
| CMO304 | RFTOC | No | Yes | Yes | ALI | $4*10^5$ | No | | | |
| CMO305 | RFTOC no MEFs | No | Yes | No | ALI | $2*10^5$ | No | | | |
| CMO306 | Sorted RFTOC | Yes | Yes | Yes | ALI | $4*10^5$ | No | $1*10^5$ | $4*10^4$ | $5*10^4$ |
| CMO307 | Unsorted mTO | No | Yes | No | Gri3D | $2*10^5$ | Yes | | | |
| CMO308 | High FTM mTO (1) | Yes | Yes | No | Gri3D | $1.9*10^5$ | Yes | $1*10^5$ | $4*10^4$ | $5*10^4$ |
| CMO309 | High FTM mTO (2) | Yes | Yes | No | Gri3D | $1.9*10^5$ | Yes | $1*10^5$ | $4*10^4$ | $5*10^4$ |
| CMO310 | High FTM mTO (3) | Yes | Yes | No | Gri3D | $1.9*10^5$ | Yes | $1*10^5$ | $4*10^4$ | $5*10^4$ |
| CMO311 | mTO- FTM (1) | Yes | No | No | Gri3D | $1.5*10^5$ | Yes | $1*10^5$ | | $5*10^4$ |
| CMO312 | mTO- FTM (2) | Yes | No | No | Gri3D | $1.5*10^5$ | Yes | $1*10^5$ | | $5*10^4$ |

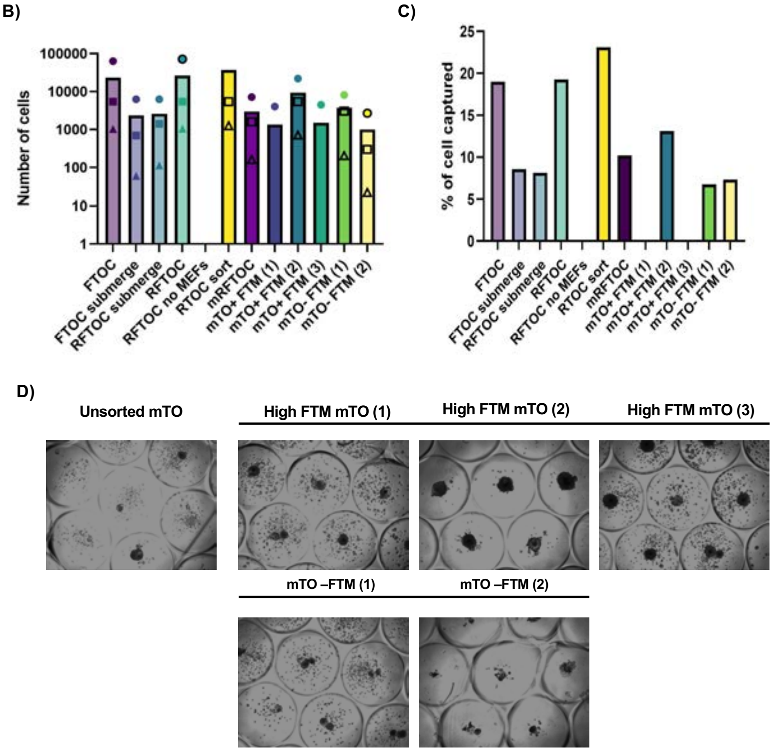


**Figure S7. Cell inputs and outputs from 'Watchbreaker' scRNAseq experiment.** (A) Table showing details of each experimental condition set up for the 10x experiment. (B, C) Graphs show (B) total number of cells counted (circle), loaded on the chip (square) and sequenced (triangle) for each condition and (C) percentage of cells captured by sequencing for each of the experimental conditions; capture efficiency calculated by the ratio of cells sequenced/ cells counted. Note that due to clumping during sample preparation the proportion of cells captured by sequencing is relatively low; this explains the lack of recovery of cells from the mTO (1) and mTO (3) conditions. (D) Images show representative microwells for each of the mTO conditions.

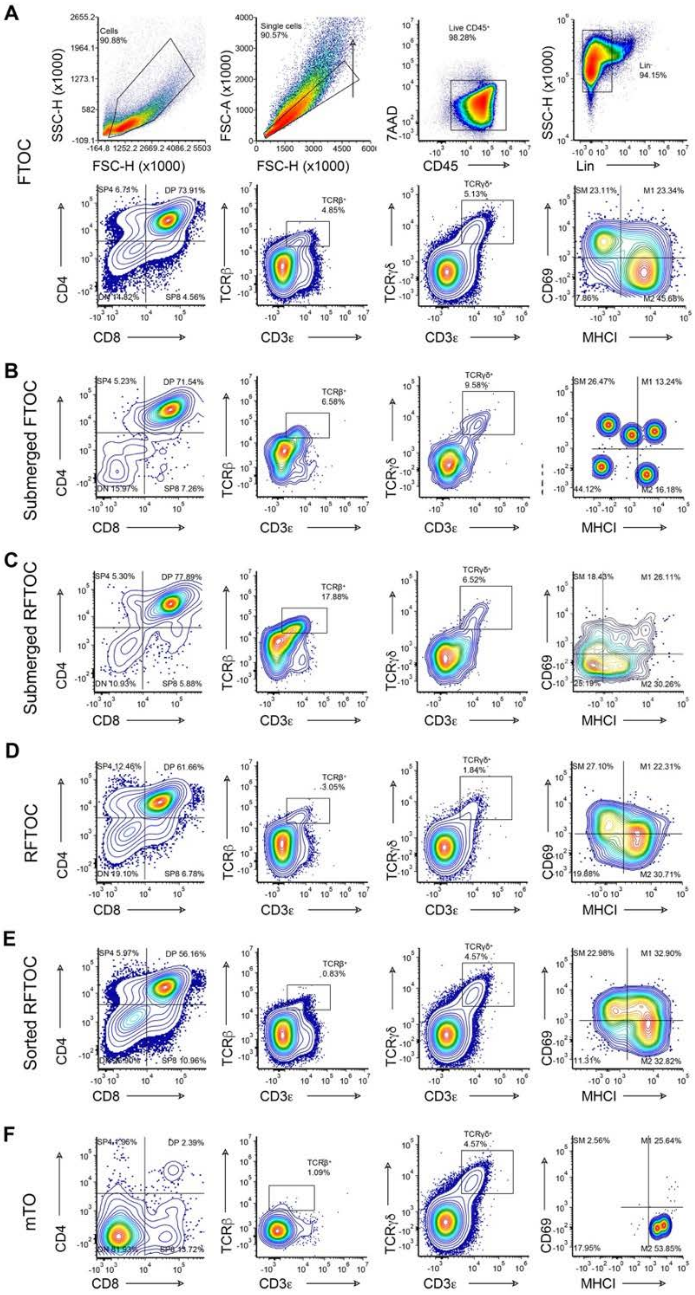


**Figure S8. Thymocyte development in different conditions in the 'Watchbreaker' 10x experiment.** Plots show thymocyte subset analysis of CD45$^{+}$ cells from the conditions shown after seven days of culture. Markers are as shown. (A) Plots show data from FTOC condition (CMO301), including gating strategy. (B-F) Plots show data from the conditions shown; Submerged FTOC (CMO302), Submerged RFTOC (CMO303), RFTOC (CMO304), Sorted RFTOC (CMO306), mTO replicate 2 (CMO309). Note that no DN to DP progression was observed in the following conditions, and therefore those data are not shown: RFTOC without MEFs (CMO305), mTO replicate 1 (CMO308), mTO replicate 3 (CMO310), mTO without FTM replicates 1 and 2 (CMO311 and CMO312). Among these, a DN population was present in all of these conditions except RFTOC without MEFs (CMO305) and the two mTO without FTM replicates.Very few cells were recovered from the unsorted mTO condition (CMO307) those data are also not shown.

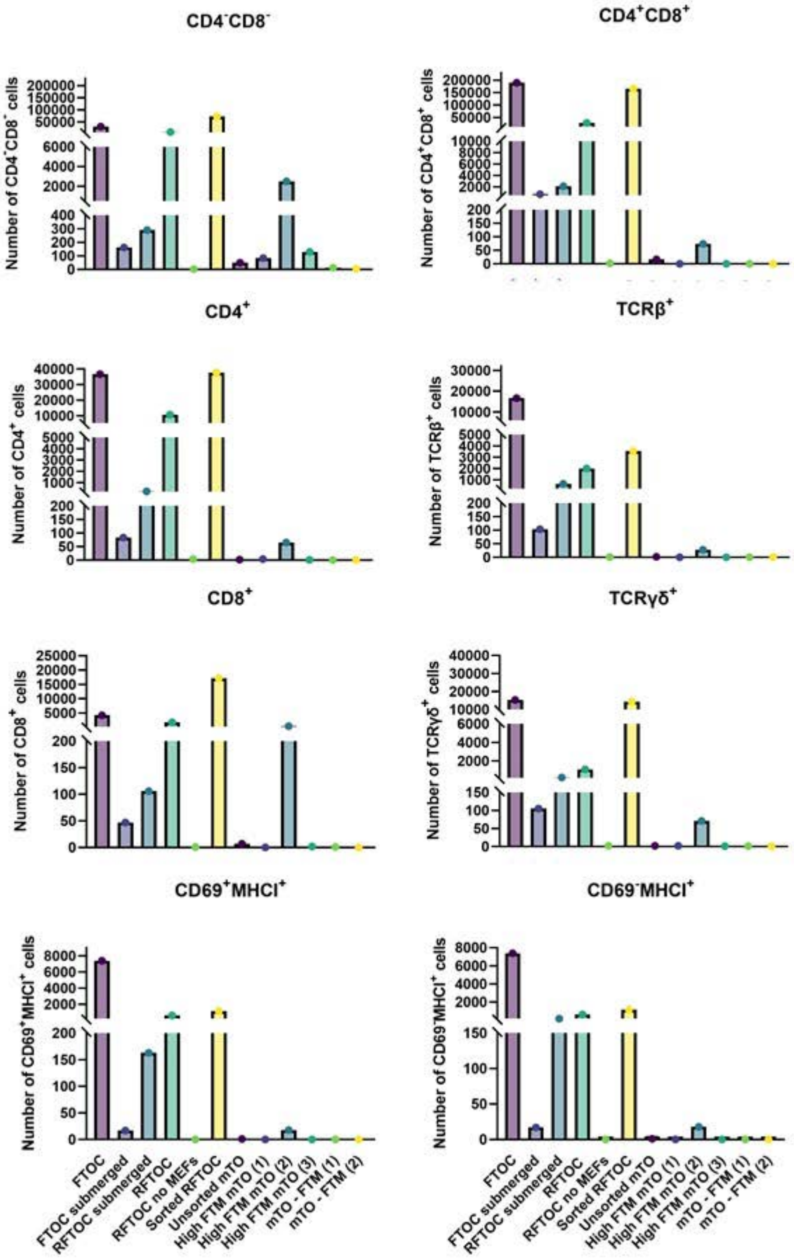


**Figure S9. Thymocyte subset numbers in each Watch-breaker condition in the 'Watchbreaker' 10x experiment.** FTOC, RFTOC and mTO were established as described in Supplemental Figure 7 and were cultured for 7 days, after which thymocyte development was assessed by analysing the $CD45^+$ population for the following subsets (see Supplemental Figure 8 for facs plots): DN ($CD4^-CD8^-$), DP ($CD4^+CD8^+$), ($CD3\varepsilon^+TCR\beta^+$), SP4 ($CD4^+$), SP8 ($CD8^+$), $CD69^+MHCI^+$, $CD69^+MHCI^-$, γδ T cells ($CD3\varepsilon^+TCR\gamma\delta^+$). A) Graphs show cell numbers recovered for each subset, for each condition. N=1 B) Graphs show percentages of parent gates present in each condition, for the subsets shown.

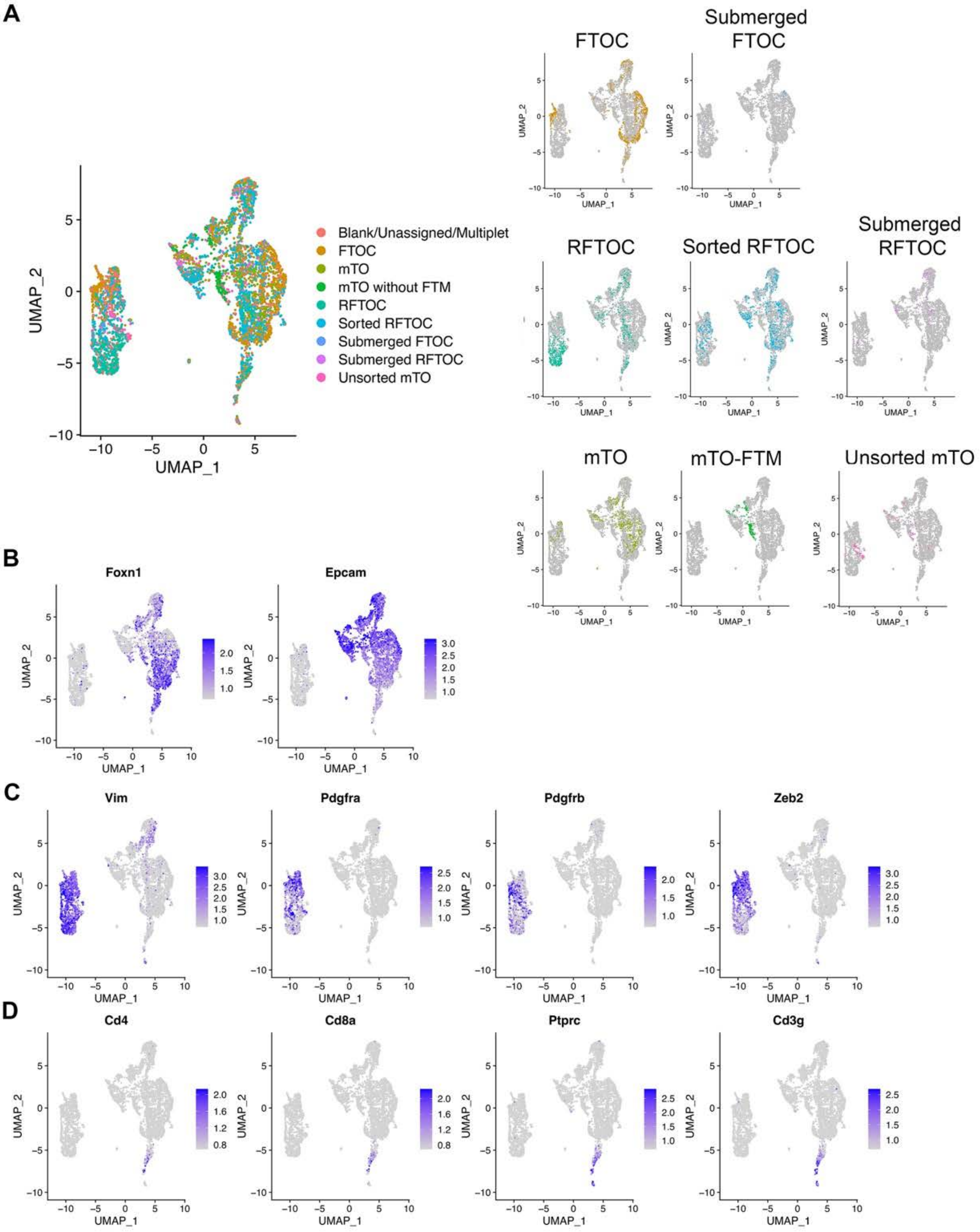


**Figure S10. Dimensional reduction showing scRNAseq data from all cells captured from all conditions.** (A) UMAP shows distribution of cells from each condition across the combined dataset. Data shown are the 4919 cells remaining after initial quality control. (B-D) Plots show distribution across combined dataset of markers for TEC (B; *Foxn1*, *Epcam*), mesenchymal cells (C; *Pdgfra*, *Pdgfrb*, *Vim, Zeb2*) and thymocytes (D; *Cd3g*, *Cd4*, *Cd8a*, *Prprc*).

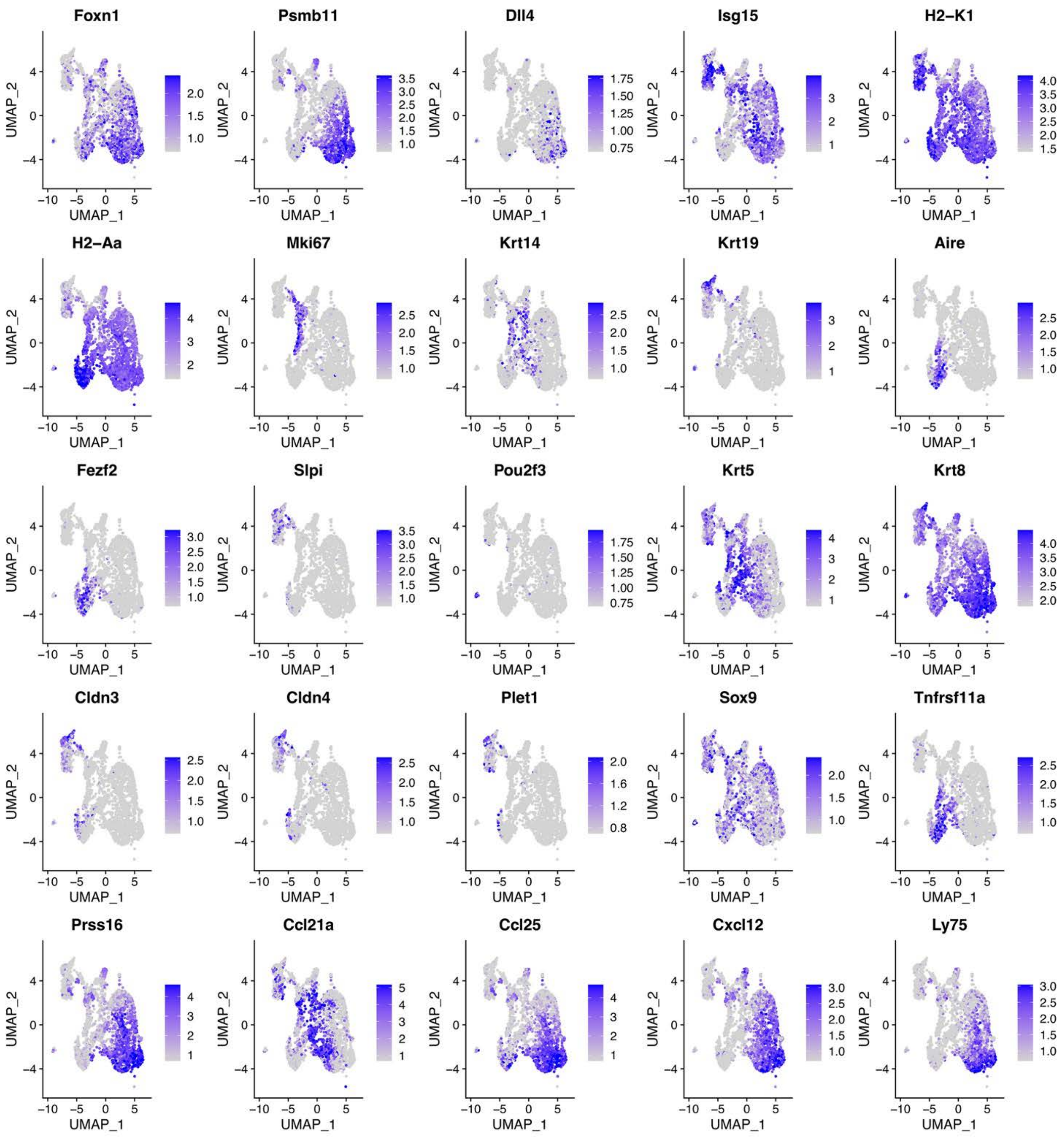


**Figure S11. Distribution of markers across TEC.** UMAPs show distribution of the markers shown across all TEC in the FTOC, RFTOC, sorted RFTOC, mTO and mTO-FTM conditions.

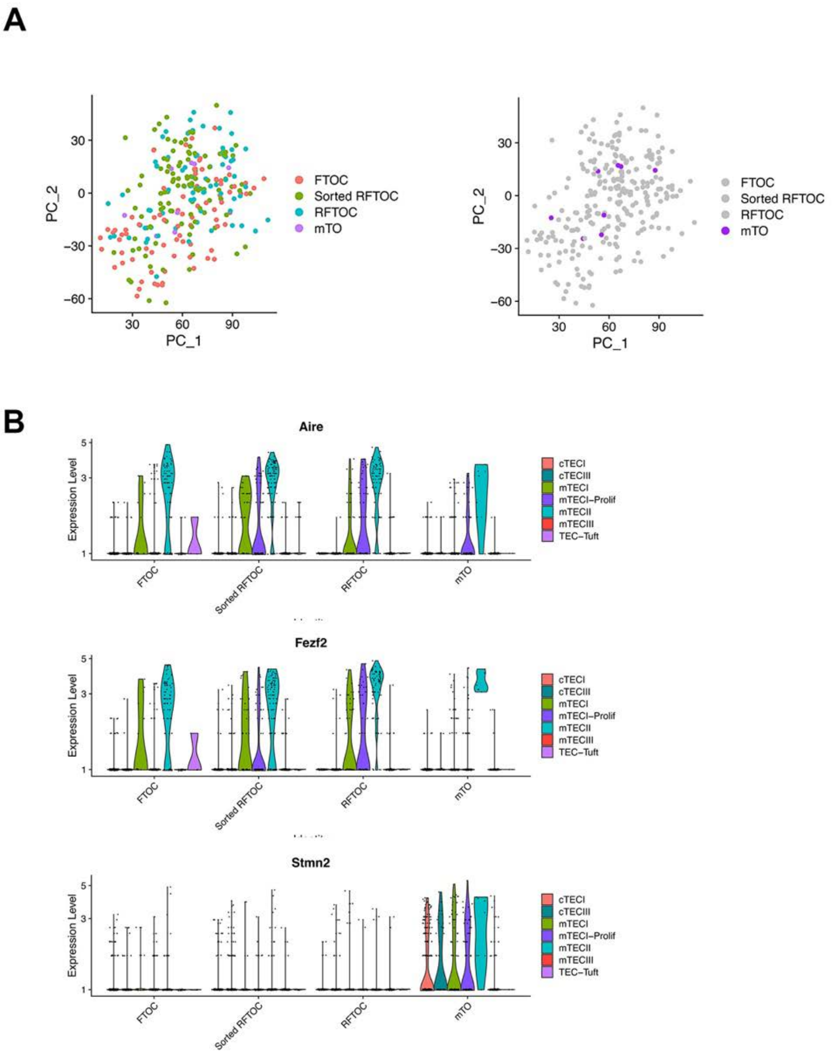


**Figure S12. Analysis of mTECII subset across four Watchbreaker conditions. (A)** PCA shows distribution of mTECII cells in FTOC, RFTOC, Sorted RFTOC and mTO. **(B)** Violin plots showing expression profiles of Aire, Fezf2 and Stmn2 in the different TEC subsets, for each of the above four conditions.

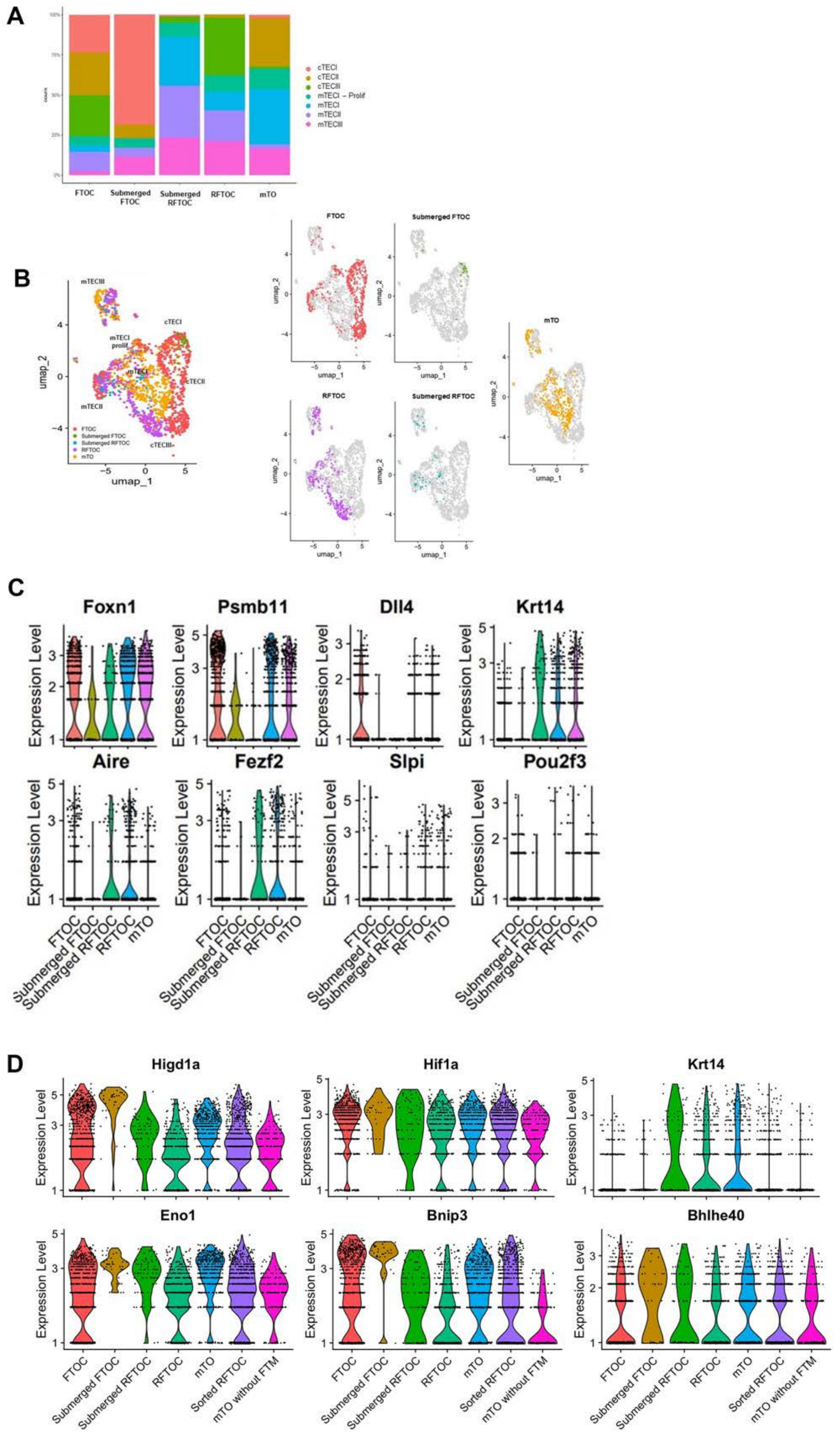


**Figure S13. Effect of submersion on cell distribution and gene expression in FTOC and RTOC. A, B)** Distribution of cell types in the FTOC, Submerged FTOC, Submerged RFTOC, RFTOC, and mTO conditions. UMAPs in (B) show combined data from the conditions shown (left panel) and distributions of TEC in individual conditions (middle and right panels). **C, D)** Violin plots show the expression profiles of the genes shown in the FTOC, Submerged FTOC, Submerged RFTOC, RFTOC, and mTO conditions. Plots in (**C**) show TEC markers; plots in (**D**) show hypoxia response genes.

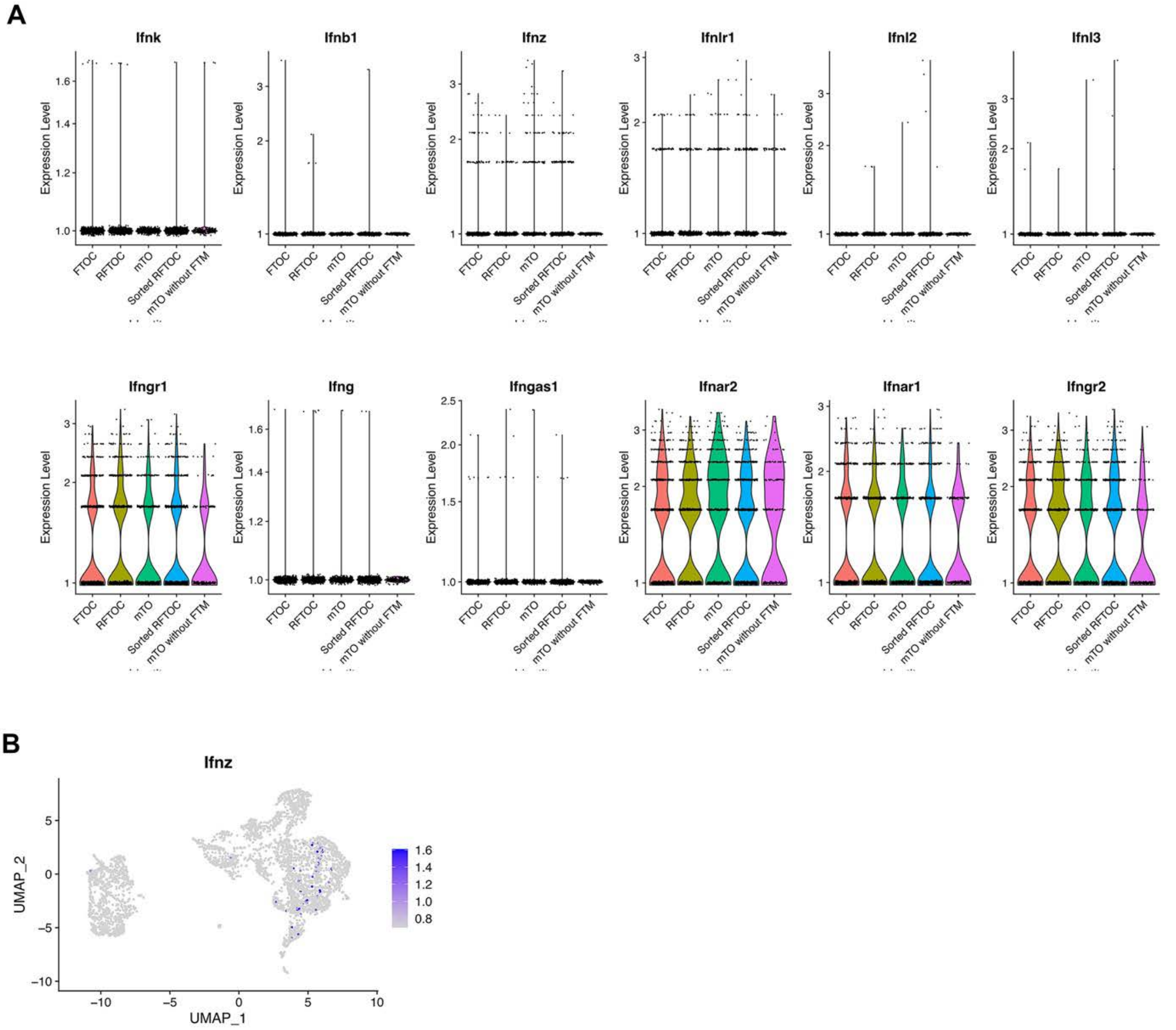


**Figure S14. Expression of IFN family and IFN receptor genes in selected 'Watch-breaker' conditions. (A)** Violin plots show the expression profiles of the genes shown in the FTOC, Sorted RFTOC, RFTOC, mTO and mTO-FTM conditions. Analyses show gene expression across all cells in the dataset, including mesenchymal cells. IFN$\alpha$ and IFN$\gamma$ receptors were expressed, but expression of the IFN family genes analysed was not detected, or was detected in only very few cells in each condition (*Ifnz*). Ifna was not detected. (B) UMAP shows distribution of *Ifnz*+ cells.

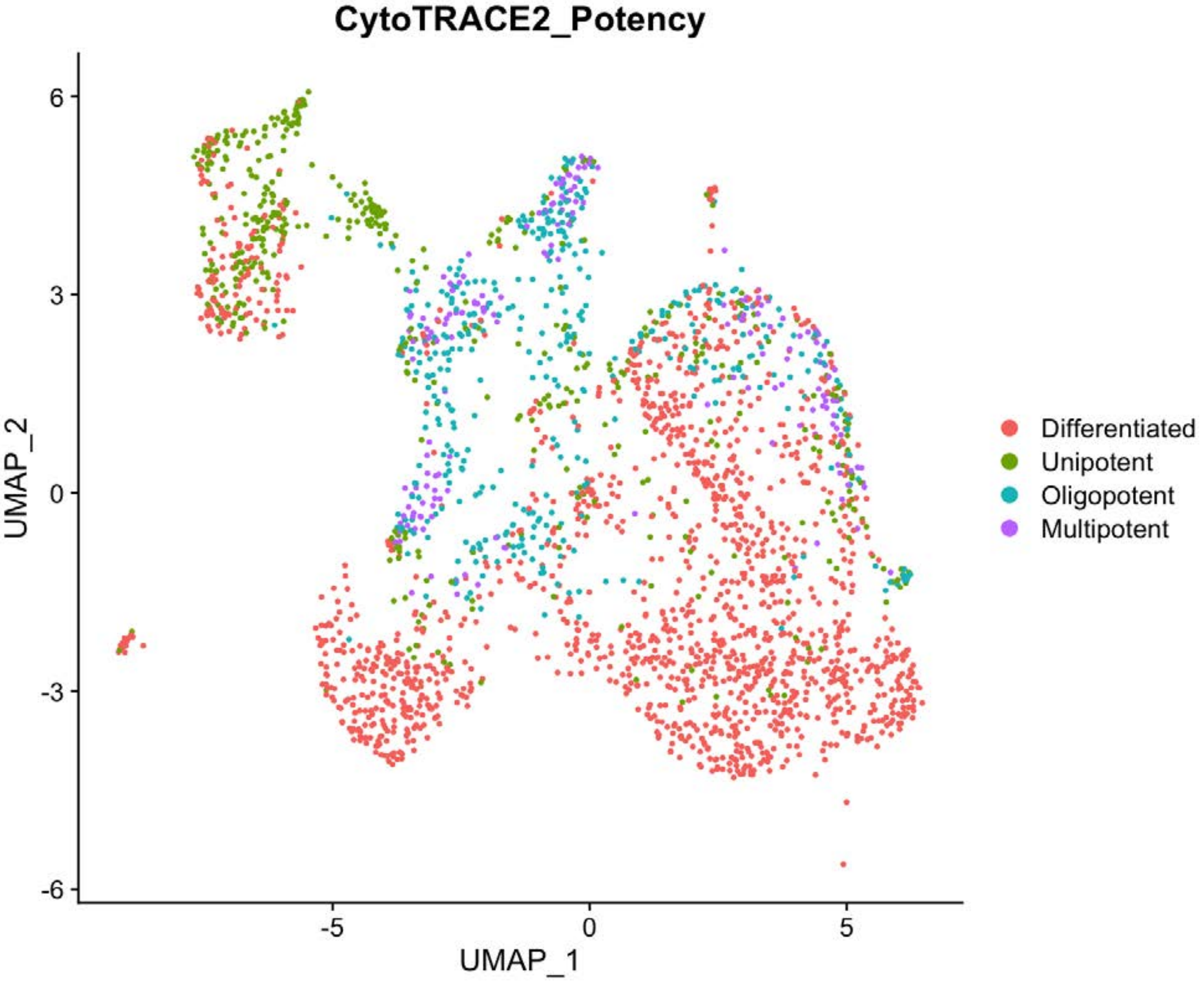


**Figure S15. CytoTRACE2 analysis across all 'Watch-breaker' conditions.** The UMAP projection is as in Figure 6. Colours indicate cell potency as predicted by the CytoTRACE2 algorithm. The mTECIII cluster is predicted to contain both unipotent progenitors and differentiated cells.

**Supplemental Tables**

| *Transgene* | *Orientation* | *Sequence* |
|---|---|---|
| Cxcl12 WT | Forward | 5’ CTG GTT TTC GCC TCT AAA GC |
| Cxcl12 Tg | Forward | 5’ TCG GCA AAA TCC CTT ATA AAT C |
| Cxcl12 | Reverse | 5’ CAG AGC TGC GAG CCT TTC |
| Rank Venus WT | Forward | 5’ CTGCGTGCTGCTCGTTCCAC |
| Rank Venus Tg | Forward | 5’ GAAGAACGGCATCAAGGCCAACTTC |
| Rank Venus | Reverse | 5’ CTGTACATACAATCTTACTGAGGGTCGC |
| GFP | Forward | 5’ TAT ATC ATG GCC GAC AAG CA |
| GFP | Reverse | 5’ GAA CTC CAG CAG GAC CAT GT |

**Table S1.** Primers used to genotype transgenic lines.

| | Antigen | Fluorophore | Dilution | Supplier | Cat. Number | Clone |
|---|---|---|---|---|---|---|
| Flow cytometric isolation of TEC, DN thymocytes, FTM and EC from E14.5 thymic lobes | EpCAM | FITC | 1:1000 | BioLegend | 118208 | G8.8 |
| | PDGFRα | APC | 1:2000 | BioLegend | 35908 | APA5 |
| | PDGFRβ | APC | 1:2000 | BioLegend | 136007 | APB5 |
| | TER119 | PE | 1:3000 | BioLegend | 116208 | TER-119 |
| | CD31 | Pe/Cy7 | 1:1600 | BioLegend | 102523 | MEC13.3 |
| | CD45 | APC-eFLuor780 | 1:1000 | eBioscience | 47-0451-82 | 30-F11 |
| | DAPI | - | 0.5 μg/ml | Life Technologies | D1306 | - |
| Flow cytometric isolation of LMPPs from E14.5 fetal liver | DAPI | - | 0.5 μg/ml | Life Technologies | D1306 | - |
| | CD5 | FITC | 1:1000 | BioLegend | 100605 | 53-7.3 |
| | CD19 | FITC | 1:500 | BioLegend | 152403 | 1D2 |
| | Gr1 | FITC | 1:800 | BioLegend | 108405 | RB6-8C5 |
| | NK1.1 | FITC | 1:800 | BD Biosciences | 553164 | PK136 |
| | CD45R/B220 | FITC | 1:800 | BioLegend | 103206 | RA3-6B2 |
| | TER-119 | FITC | 1:800 | eBioscience | 11-5921-85 | TER-119 |
| | F4/80 | FITC | 1:1600 | BioLegend | 2BM8 | 123108 |
| | CD135 (Flt3) | APC | 1:50 | BioLegend | 135310 | A2F10 |
| | Sca-1 | PE | 1:2000 | BioLegend | 108107 | D7 |
| | CD117 (cKit) | PE/Cy7 | 1:1600 | BioLegened | 105813 | 2B8 |
| Flow cytometric analysis of T cell subsets from mTOs | 7AAD | - | 1:200 | BioLegend | 420403 | - |
| | CD11b | FITC | 1:800 | BioLegend | 101205 | M1/70 |
| | CD11c | FITC | 1:800 | BioLegend | 117305 | N418 |
| | Gr1 | FITC | 1:800 | BioLegend | 108405 | RB6-8C5 |
| | NK1.1 | FITC | 1:800 | BD Biosciences | 553164 | PK136 |
| | CD45R/B220 | FITC | 1:800 | BioLegend | 103206 | RA3-6B2 |
| | TER-119 | FITC | 1:800 | eBioscience | 11-5921-85 | TER-119 |
| | EpCAM | FITC | 1:800 | BioLegend | 118208 | G8.8 |
| | CD4 | PE | 1:800 | BioLegend | 100512 | RM4-5 |
| | CD8α | APC | 1:400 | eBioscience | 17-0081-82 | 53-6.7 |
| | CD3ε | BV785 | 1:100 | BioLegened | 100355 | 145-2C11 |
| | TCRβ chain | PE/Cy7 | 1:100 | BioLegend | 109222 | H57-597 |
| | CD45 | APC/eFlour 780 | 1:800 | eBioscience | 47-0451-82 | 30-F11 |
| | γ/δTCR | BV605 | 1:400 | BioLegend | 118124 | GL3 |
| | MHC1 (H-2k) | BV510 | 1:400 | BioLegend | 116523 | AF6-88.5 |

| | | | | | | |
|---|---|---|---|---|---|---|
| | CD69 | BV421 | 1:100 | BioLegend | 104528 | H1.2F3 |
| IHC of in vitro cultures. | Vimentin | - | 1:250 | Abcam | ab92547 | Abcam (rabbit) |
| | MHCII | - | 1:200 | Abcam | ab15630 | ER-TR3 |
| | β5t | - | 1:100 | MBL | PD021 | Polyclonal |
| | DLL4 | - | 1:50 | BioLegend | 130802 | HMD4-1 |
| | MHCI | - | 1:200 | BDBioscience | 550550 | AF6-88.5 |
| | HOECHST | - | 1:1000 | Life Technologies | 62249 | |
| | Anti-Rabbit secondary | Alexa Fluor 568 | 1:1000 | Invitrogen | A-11011 | |
| | Anti-Rat secondary | Alexa Fluor 647 | 1:1000 | Invitrogen | A-21247 | |

**Table S2.** Antibodies used in panels for flow cytometry and immunohistochemistry. Table shows the name, associated fluorophore where appropriate, supplier and clone number. DAPI, (4',6-Diamidino-2-Phenylindole, Dihydrochloride); IHC, immunohistochemistry.

| Antibody | Barcoded Reagent | Dilution | Supplier | Cat. Number |
|---|---|---|---|---|
| UEA1-Biotin | TotalSeqTM-SAV-Pe- B0952 | 1:1500 | Vector Laboratories | B-1065 |
| MHCII (α-mouse I-A/I-E) | TotalSeq™ - B0117 | 1:10000 | BioLegend | 107657 |
| CD40 | TotalSeq™ - B0903 | 1:3200 | BioLegend | 124639 |
| CD80 | TotalSeq™ - B0849 | 1:3200 | BioLegend | 104757 |
| EpCAM | TotalSeq™ - B0449 | 1:800 | BioLegend | 118247 |
| CD45 | Beads | 1:10 | Miltenyi Biotec Inc. | 130-052-301 |

**Table S3.** Cell staining reagents used in 10x sequencing experiment.

**Table S4.** List of genes differentially expressed between mTO and FTOC, mTO and RFTOC and mTEC and Sorted RFTOC, across all TEC for each condition. Positive fold change (FC) indicates upregulated in mTO. Uploaded separately.

**Table S5.** List of genes differentially expressed in mTECII cells, between mTO and FTOC, mTO and RFTOC and mTEC and Sorted RFTOC. Positive fold change (FC) indicates upregulated in mTO. Uploaded separately.

**Table S6.** List of genes differentially expressed between mTO+FTM and mTO-FTM across all TEC for each condition. Positive fold change (FC) indicates upregulated in mTO+FTM. Uploaded separately.

**Table S7.** List of genes differentially expressed between mTO+FTM and mTO-FTM for cTECI/cTEC-neg and mTECIII cells. Positive fold change (FC) indicates upregulated in mTO+FTM. Uploaded separately.
